# A cellular and spatial map of the choroid plexus across brain ventricles and ages

**DOI:** 10.1101/627539

**Authors:** Neil Dani, Rebecca H. Herbst, Naomi Habib, Joshua Head, Danielle Dionne, Lan Nguyen, Cristin McCabe, Jin Cui, Frederick B. Shipley, Ahram Jang, Christopher Rodman, Samantha J. Riesenfeld, Feng Zhang, Orit Rozenblatt-Rosen, Aviv Regev, Maria K. Lehtinen

## Abstract

The choroid plexus (ChP), located in each brain ventricle, produces cerebrospinal fluid (CSF) and forms the blood-CSF barrier, but is under-characterized. Here, we combine single cell RNA-Seq and spatial mapping of RNA and proteins to construct an atlas of each ChP in the developing and adult mouse brain. Each ChP comprises of epithelial, endothelial, mesenchymal, immune, neuronal, and glial cells, with distinct subtypes, differentiation states and anatomical locations. Epithelial, fibroblast, and macrophage populations had ventricle-specific, regionalized gene expression programs across the developing brain. Key cell types are retained in adult, with loss of developmental signatures and maturation of ventricle-specific regionalization in the epithelial cells. Expression of cognate ligand-receptor pairs across cell subtypes suggests substantial cell-cell interactions within the ChP. Our atlas sheds new light on the development and function of the ChP brain barrier system, and will facilitate future studies on its role in brain development, homeostasis and disease.

## INTRODUCTION

The choroid plexus (ChP) forms a blood-cerebrospinal fluid (CSF) barrier in each brain ventricle and is essential for the development and function of the brain. Built as an epithelial bilayer with an accompanying network of predominantly non-neural cell types and vasculature, the ChP regulates the secretion and composition of CSF that fills the brain’s ventricles, bathes stem cells, and reaches neurons via exchange with the interstitial fluid^1^. CSF composition is regulated by *de novo* synthesis and secretion of signals by epithelial cells, as well as selective transcytosis of blood-borne factors^2–5^, which regulate neurogenesis^6,7^ or guide migration of newborn neurons^8^. The ChP is also sensitive to peripheral body signals, and arousal states ^9,10^ and gates immune cell passage from body to brain^3,11–13^. Disrupted ChP function and CSF composition, as well as abnormal CSF volume and ventricle space, are common to neurologic disease, including brain inflammation^14^, autism^15,16^, schizophrenia^17^, and Alzheimer’s disease^18,19^.

Despite these essential roles, remarkably little is known regarding the molecular mechanisms governing these functions of the choroid plexus, its cellular networks, and histology. In particular, our knowledge of the cellular composition of the choroid plexus within each ventricle in developing and adult brain is limited. Each choroid plexus develops independently from distinct locations along the roofplate, where capillaries, mesenchymal and neural crest cells invaginate the neuroepithelium^1,20,21^. Bulk profiling of whole tissue^4,7,22^ or sorted subpopulations^4^ indicates that each ChP retains a developmental blueprint of its origins along the body axis, with ventricle-specific transcriptomes and secretomes for the lateral and 4th ventricle ChPs^4^. The less-accessible third ventricle remains even less explored, and its boundaries with the adjacent neural tissues remain undefined. Earlier efforts to address these deficiencies were limited by the lack of techniques to access, isolate and comprehensively characterize ChPs, thus limiting our ability to understand their functions and harness them for therapeutic benefit.

Here, we characterized the choroid plexus across all brain ventricles by single cell RNA-Seq and spatial mapping of specific RNAs and proteins, in the developing and adult mouse brain. We profiled 15,620 cells and 62,474 nuclei, followed by a multi-tiered validation with fluorescent *in situ* hybridization (FISH), immunostaining, viral labeling, and transgenic lines. We employed whole tissue imaging to map the identified cell types, sub-types and states to uncover specialized niches within ChPs and along transition areas between the ChP and adjacent brain. We charted ventricle-specific diversity in ChP epithelial, fibroblast and immune cells in the developing brain and compared these features to the adult. Cells expressed distinct ligands and receptors, suggesting functional interaction networks within the ChP with likely roles in maintaining blood-CSF barrier integrity and CSF secretion. Our molecularly-guided approach to spatially map the ChPs provides a resource for creating tools to access and control this essential brain-body barrier.

## RESULTS

### Molecular survey defines the cellular composition of the developing ChP across ventricles

To chart a cell atlas of the developing ChP, we profiled 15,620 single cells from the ChP of each brain ventricle (lateral, third and fourth ventricles; LV, 3V, 4V) at embryonic day [E]16.5 (Fig. 1A-C). We first refined microdissection techniques to isolate the ChPs in the developing mouse brain (Fig. 1A) and optimized tissue dissociation. Next, we collected healthy cells by fluorescence-activated cell sorting (FACS) across three independent experiments and nine litters of mice and profiled them by droplet-based single cell RNA-seq (scRNA-Seq, **Methods**, Fig. 1B, S1A). We partitioned the cells into clusters (**Methods**) followed by *post hoc* annotation by expression of canonical cell markers, which identified epithelial, mesenchymal (mural and fibroblast), endothelial, immune, neuron and glia-like cell types in each ChP from all ventricles (Fig. 1C,D, S1B), and associated each cell type with marker genes (Fig. 1D). We used canonical markers to determine the spatial positions of each major cell type within the ChP tissue from each ventricle (Fig. 1E, S1C). Notably, actively cycling cell subsets were present within each of the six major cell classes (Fig. S1D), and positionally enriched along the base of the ChP proximal to the brain based on staining for the proliferation marker Ki67 (Fig. S1E).

**Fig. 1.**
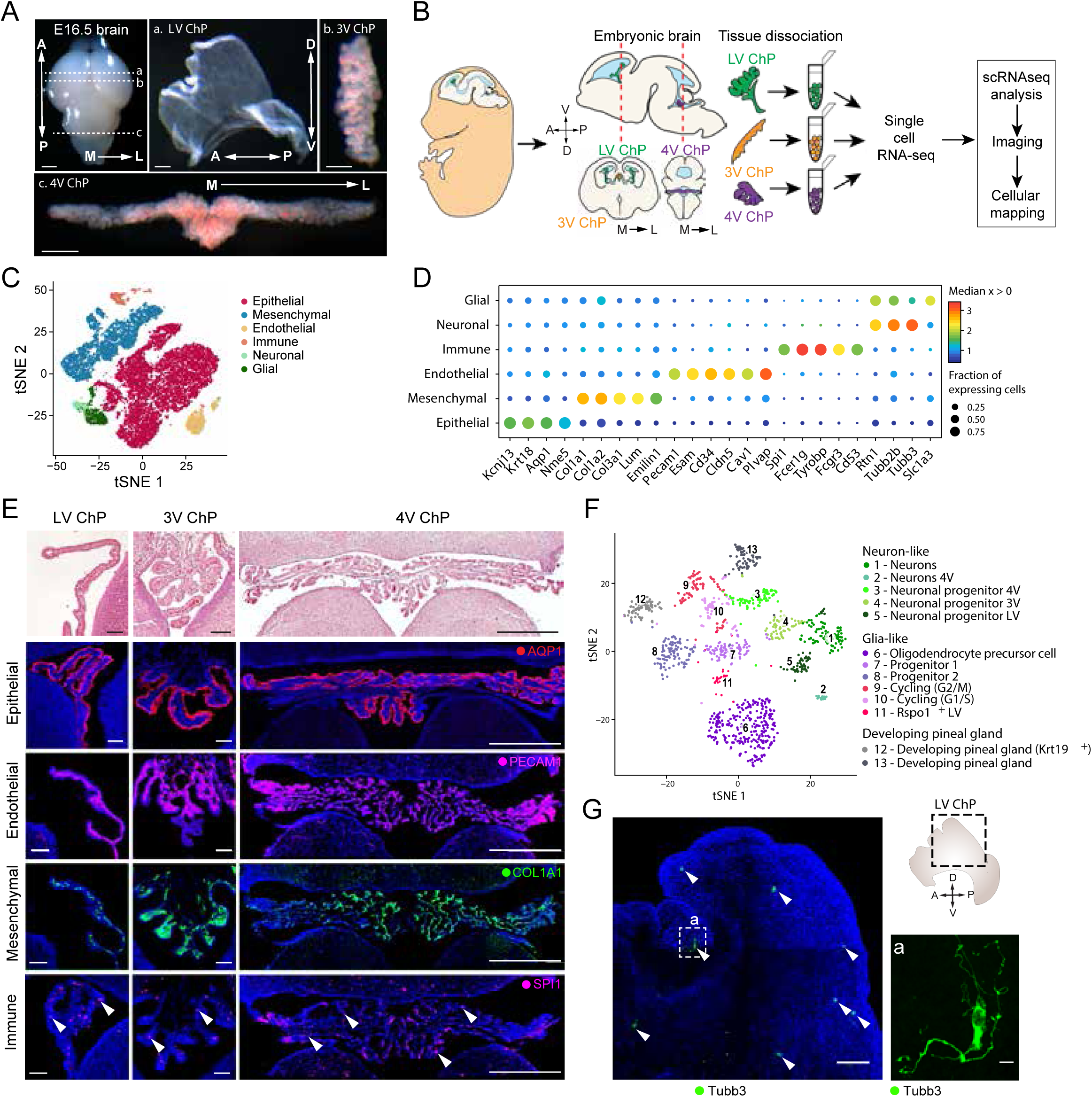
Single cell RNA-Seq of the cellular composition of ChPs. (**A**) ChP tissues from each brain ventricle. Location of ChP (horizontal dotted lines) in the E16.5 brain (top left panel, scale bar = 1 mm). Representative images of intact LV ChP (a. top middle panel, scale bar = 0.2 mm), 3V ChP (b. top right panel, scale bar = 0.2 mm) and 4V ChP (c. bottom panel, scale bar = 0.5 mm). Arrows: anterior-posterior (A/P), dorso-ventral (D/V) and medial-lateral (M/L) axes. **(B)** Workflow. **(C)** Major cell subsets of the embryonic ChP. A 2D t-stochastic neighborhood embedding (t-SNE) of 15,620 single cell profiles (n=9 mice, in pools of 3 animals per ventricle), colored by *post hoc* annotated cell type. **(D)** Canonical and novel cell type markers. Median expression level in expressing cells (color) and proportion of expressing cells (circle size) of selected genes (columns) in each major cell population (rows). **(E)** Immunostaining of major cell type markers reveal shared features of each ChP. Top panels: Coronal sections of E16.5 brain show LV (left), 3V (middle) and 4V (right) ChP in cross-section (H&E stained). Lower panels: Immunostaining of major cell type markers in the superficial epithelial cell layer (AQP1), sub-epithelial stromal space that contains mesenchymal (COL1A1), endothelial (PECAM1) and immune cells (SPI1, white arrowheads). Scale bar (LV ChP and 3V ChP) = 20 µm, scale bar (4V ChP) = 500 µm. **(F)** Neuronal and glia-like cell subsets. tSNE of neuronal and glia-like cell profiles, colored and numbered by cluster membership. **(G)** Neuronal cell bodies in the ChP. Whole explant imaging of LV ChP stained with TUBB3 antibodies (green). White arrowheads: neuronal cell bodies within the plexus (Scale bar = 100 µm), Inset a: Neuronal (TUBB3) cell morphology (scale bar = 10 µm). Schematic with dotted outline (black): region of LV ChP depicted in immunostaining. Double headed arrows: A/P and D/V axes.

### Neurogenic and gliogenic cell populations are found within all developing ChPs

In agreement with previous studies documenting the presence of neuronal cell bodies and neural innervation of the ChP^23^, we captured both neuronal (*Tubb3* expressing) and glial-like (*Slc1a3*/EAAT1 expressing) populations in each developing ChP (Fig. S1F,G) in all developing ChPs (Fig. S1B). Subsets of glial-like cells expressed markers of glia-neuron progenitors/stem-like cells (*Rspo2/3, Nes*, *Sox2*, *Fabp7, Hes1, Pax6*) as well as oligodendrocyte precursor cells (OPCs, *Olig1, Olig2*)^24^ (Fig. 1F, S1H). Subsets of neurons expressed markers of immediate progenitor (*Eomes/Tbr2*) and immature neurons (*Neurod1/2, Dlx1/2*, Fig. 1F, S1H), which also expressed a range of neuropeptides (Fig. S1H). We confirmed the presence of differentiated neurons using whole LV ChP explants (**Methods**, Fig. 1G), leveraging the relatively simpler three-dimensional structure of LV ChP (Fig. 1A) to facilitate whole tissue imaging, reconstruction and cell identification. Moreover, some of the neuronal cell bodies with processes within the LV ChP stained positive for the neurotransmitter serotonin (5-HT) (Fig. S1I). This may be related to earlier findings in the adult brain that serotonergic axons from brain regions such as the raphe nucleus innervate the ChP^25^. Finally, some of these ChP neurons express *Syn1*, another marker of neuronal identity (Fig. S1J), and were correspondingly amenable to adeno-associated viral transduction (AAV1.Syn1, Fig. S1I), emphasizing the potential for their transgenic perturbations.

Two subsets in the 3V expressed markers of developing pinealocytes (*e.g., Crx*, *Krt19*; Fig.s 1F, clusters 12 and 13, Fig. S1F,H). The pineal gland develops caudal to the 3V ChP, and these two secretory tissues form together the roof of the 3V^26^. We leveraged the inclusion of ChP adjacent brain tissue in our cell atlas to molecularly map the boundary between the 3V ChP and developing pineal gland (Fig. S1K).

### An inferred differentiation continuum suggests a common progenitor of epithelial and neural cells

Surprisingly, the epithelial and neuronal / glia-like cells were not discretely separable (Fig. 1C) by any low-dimensional embedding analyses that we attempted (**Methods**), and stem cell marker genes were expressed in cells at their intersection point (Fig. S2A), suggesting a model where a common progenitor pool for both epithelial and neuronal cells may exist in the ChP. To explore this hypothesis, we modeled the predicted phenotypic continuum between neuronal, glia-like and epithelial cells using a diffusion map^27^ of cells of the 3V ChP. The diffusion embedding arranged neurons and mature epithelial cells at two opposite ends of a trajectory, with a progenitor glia-like population (*Rspo2+*) between them (Fig. 2A). *Rspo2^+^* progenitor like cells also expressed several stem markers (Fig. S2B,C, progenitor 1 and 2 populations in Fig. 1F; **Methods**), which have previously been shown in precursor cells of the developing cortex that have glia like properties and give rise to ependymal cells (modified epithelial cells) and neurons^24^.

**Fig. 2.**
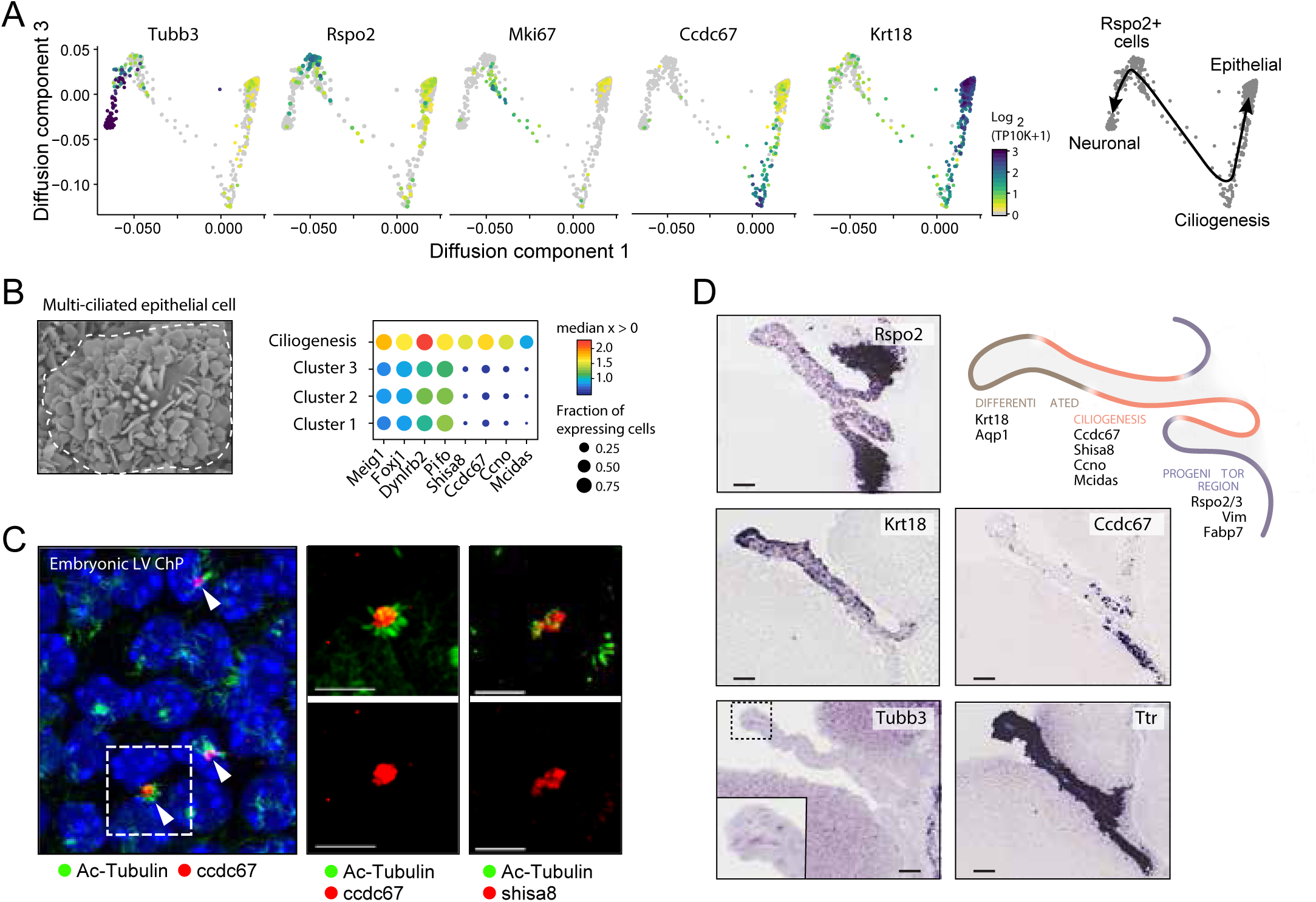
An epithelial differentiation trajectory suggests a common progenitor of epithelial and neural cells. (**A)** Inferred differentiation trajectory from common progenitor to epithelial and neuronal cells. Diffusion map 2-D embedding of neuronal, glia-like and epithelial cell profiles (dots) from 3V ChP, displaying diffusion components 1 (x axis) and 3 (y axis), colored by log_2_(TP10K+1) expression of marker genes of (from left to right): neuronal (*Tubb3*), progenitors (*Rspo2*), cycling (*Mki67*), ciliogenesis (*Ccdc67*), and mature epithelial (*Krt18*) cells. Right: Schematic model of suggested differentiation trajectories. **(B)** Multi-ciliated ChP epithelial cells. Left: Scanning electron microscope image of a multi-ciliated epithelial cell from a mouse ChP (Scale bar = 1 µm). Right: Median expression level in expressing cells (color) and proportion of expressing cells (circle size) of markers of ciliogenesis and primary cilium (columns) across subsets of epithelial cells (rows, as in Fig. S2E). **(C)** Ciliogenesis in a subset of epithelial cells. Left: Whole explant imaging of the LV ChP stained with anti-Ac-Tubulin (green) and anti-CCDC67/DEUP1 (red). Right: Images of individual ciliary tufts (green) stained with anti-CCDC67 (red, middle) or anti-SHISA8 (red, right). Scale bar = 10 µm (left) or 5 µm (all other panels). **(D)** Spatial mapping of the differentiation trajectory in the LV ChP. *In situ* hybridization images from the LV ChP from Genepaint of markers of (from top): progenitors (*Rspo2*); differentiated epithelial cells (*Krt18*) ciliogenesis/basal body biogenesis (*Ccdc67*), neurons (*Tubb3*) and epithelial cells (*Ttr*) (Scale bar = 100 µm). Top right: Model of proximal to distal organization of progenitor and differentiation states in the LV ChP. Progenitors at the base LV ChP and adjacent brain, ciliogenesis/basal body biogenesis epithelial cells along the root of the LV ChP, followed by mature epithelial cells.

Our analysis also highlighted dividing (Fig. S2D) and newly differentiated post-mitotic epithelial cells that transiently expressed ciliogenesis genes (Fig. 2A, 2B, S2E), such as *Ccdc67/Deup1*, which drives centriole biogenesis, an essential step of multi-ciliation^28^. Another marker, *Shisa8*, was expressed near the base of acetylated-tubulin tufts in multi-ciliated epithelial cells (Fig. 2C), and was recently identified as a regulator in iPSC reprogramming^29^.

These populations had distinct spatial boundaries: progenitor cells were located near the ChP-brain boundary (by ISH of *Rspo2*; Fig. 2D); newly differentiated epithelial cells localized near the base of the LV ChP proximal to the brain (by ISH of *Ccdc67/Deup1*; Fig. 2D), whereas more mature epithelial cells were positioned along the length of the ChP, extending distally into the LV (ISH by *Krt18*) (Fig. 2D). This maturation gradient of ChP epithelial cells matches models based on classic electron microscopy^22,30^ and provide a molecular handle for investigating these distinct stages of epithelial cell development.

### Epithelial and mesenchymal cell programs are regionalized across brain ventricles

Epithelial cells formed the largest cell class in each ChP and partitioned into several clusters representing cycling, newly differentiated cells undergoing ciliogenesis (above), and mature cells with ventricle-specific differences (Fig. 3A). On average, the scRNA-seq data recapitulated differential gene expression between epithelial cells from LV and 4V ChP from bulk RNA-seq^4^ (Fig. S3A).

**Fig. 3.**
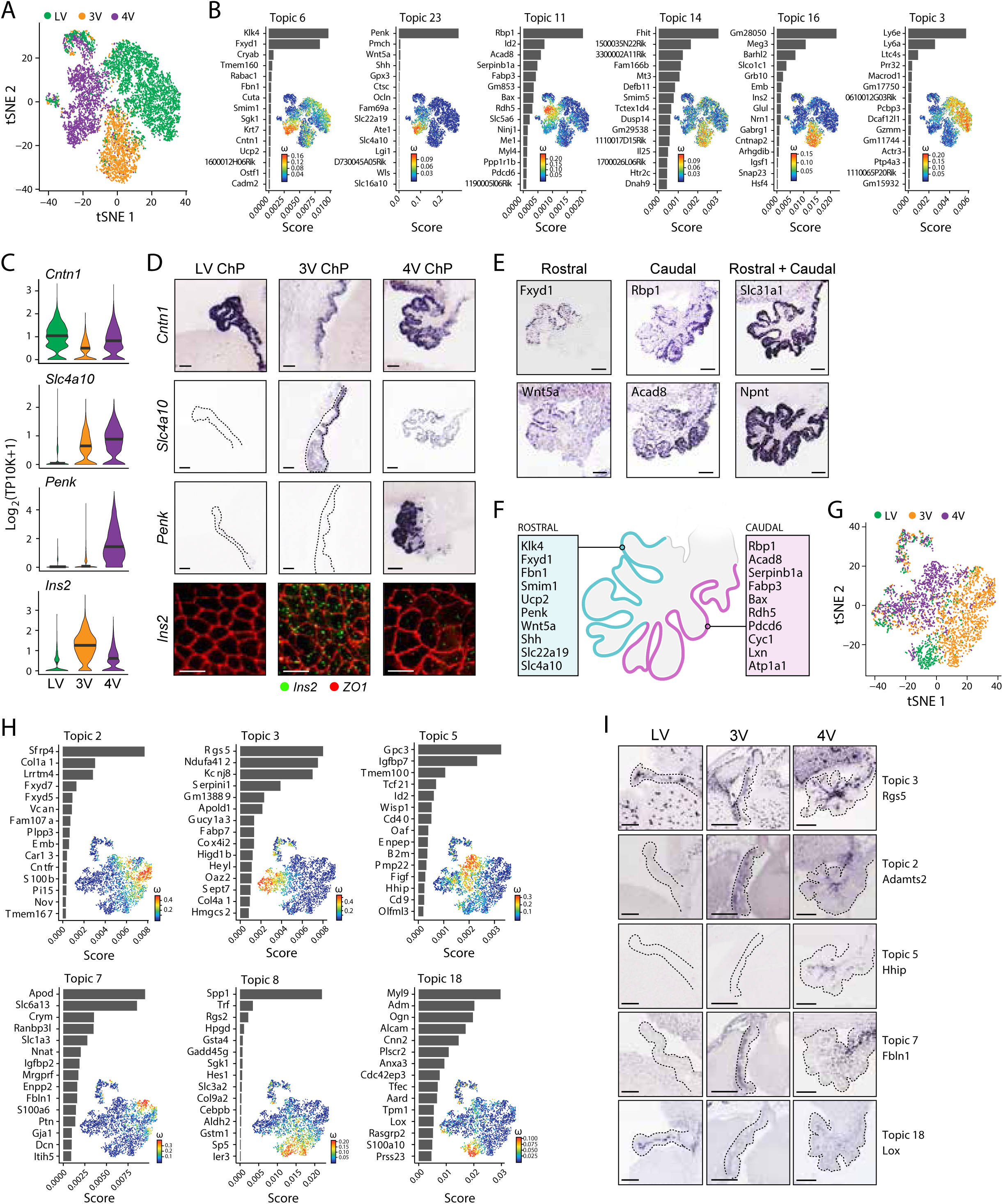
Regionalized epithelial and mesenchymal transcriptional programs across ventricles. **(A)** Distinct epithelial cells clusters by ventricle. t-SNE of epithelial cell profiles (dots), colored by ventricle. **(B)** Ventricle associated transcriptional programs in epithelial cells. For each topic that is differentially weighted between ventricles, shown is a bar plot of topic scores for top ranked genes (left), and tSNE of the cell profiles (as in A) colored by topic’s weight per cell (right). Bold: Genes highlighted in (D,E). **(C-D)** Key genes with regionalized expression in epithelial cells across ventricles. **(C)** Distribution of log_2_(TP10K+1) expression (y axis) of each denoted gene across ventricles (x axis). **(D)** Top three panels: *In situ* hybridization from Genepaint of sagittal sections of the LV, 3V and 4V ChP (columns) of genes with regionalized expression (rows) (Scale bar = 100 µm). Bottom panels: smFISH of *Ins2* in whole explants (Scale bar = 10 µm). **(E)** Regionalized expression within the 4V ChP epithelium. *In situ* hybridization from Genepaint of transcripts in sagittal sections of the 4V ChP (Scale bar = 100 µm). **(F)** Rostral-caudal patterning within the 4V ChP epithelium. Model of regionalized expression in the medial core of the 4V ChP that lies within the 4^th^ ventricle, with example genes identified by topic models and validated *in situ* (in **E**). **(G)** Mesenchymal cells largely cluster by ventricle. t-SNE of mesenchymal cell profiles (dots) colored by ventricle. **(H)** Ventricle associated transcriptional programs in mesenchymal cells. Each topic that is highly weighted in a significant fraction of mesenchymal cells (except cycling cells; Fig. S3H) is shown as in B, with the tSNE in **G**. Bold: genes highlighted in (I). **(I)** Regionalized mesenchymal transcriptional programs. *In situ* hybridization in sagittal sections of the LV, 3V and 4V ChP (Genepaint) for a pericyte marker (*Rgs5*) and genes with ventricle specific enrichment in fibroblast in the 3V (*Adamts2, Fbln1*), 4V (*Hhip*), and LV (*Lox*) (Scale bar for LV = 50 µm; 3V and 4V = 10 µm).

To identify the transcriptional programs that drive the distinction of differentiated ChP epithelial cells by brain ventricle, we applied topic modeling using Latent Dirichlet Allocation, originally introduced in natural language processing^31^ and population genetics^32^ and recently applied to scRNA-seq data^33,34^. In topic modeling, a cell is modeled as a mixture of a small number of transcriptional programs (“topics”), where each topic is a distribution over genes. A gene can belong to multiple topics with different weights, reflecting the gene’s role in each topic. Likewise, a topic’s weight for a given cell reflects the relative prominence of the corresponding biological process associated with that topic in that cell. We learned topic models for all epithelial cells, and then searched for topics that were differentially weighted across subsets of individual cells or that described an interpretable biological process, based on the associated genes (Fig. 3B, S3B,C, **Methods**), such as ciliogenesis (Topic 24, Fig. S3B, 2D), or immediate early genes (Topic 19, which may reflect a response to tissue dissociation^35^, as it was absent from single nucleus RNA-seq (snRNA-Seq, Fig. S3B,D)).

Several topics (3,6,11,14,16,23) were differentially weighted across cells from the different brain regions (Fig. S3C), including genes encoding transporters (Fig. S3E). We validated the ventricle specific enrichment of key associated genes using single molecule fluorescence *in situ* hybridization (smFISH) and published data^36^ (Fig. 3C,D). Notably, *Ins2*, encoding an insulin precursor and associated with Topic 16, was over-expressed in 3V epithelial cells (Fig. 3B-D), suggesting the 3V ChP as an internal source of insulin within the developing brain. Moreover, epithelial cells within the 4V ChP could be partitioned into two subpopulations by their weights of topic 6, 11 or 23 (Fig. 3B, E, S3F). Mapping the expression of highly scoring genes for these two topics within the 4V ChP identified a rostro-caudal gradient of gene expression along the medial core of the plexus within the fourth ventricle (Fig. 3E,F), while a subset of lowly scoring genes were expressed along the whole 4V ChP (Fig. 3E). These gradients may derive from the earliest stages of development, when roof plate progenitors originating from distinct rhombomeres give rise to hindbrain and 4V ChP^20^.

We used similar analysis to characterize regionalized features in the large and heterogeneous population of mesenchymal cells in the ChP. Mesenchymal cells partitioned into fibroblasts and mural cells (including pericytes and smooth muscle actin [SMA] positive cells) (Fig. S3G), consistent with cranial mesenchyme and neural crest contributions to the stromal space^21,37^. While pericytes (Topic 3) and proliferating mesenchymal cells (Topic 12 and 16) were similar in each ChP, topic modeling identified transcriptional programs of fibroblasts significantly enriched in specific ventricles; LV (Topic 8 and 18), 3V (Topic 2 and 7) and 4V (Topic 5) (Fig. 3G,H, S3H,I), which we validated using published data (Fig. 3I^36^). These topics revealed ventricle-dependent expression of genes encoding growth factors (*Bmp4/7, Wnt4*/*2)* and extracellular matrix proteins (Fig. 3H, S3J). For example, 4V ChP fibroblasts expressed high levels of genes encoding signaling molecules critical for hindbrain development (*e.g.*, *Hhip*^38^, *Ptch1*^39^, *Rbp4*^40^, and *Wisp1*, Fig. 3H, I)). More generally, genes involved in regulation of cell migration and tissue development were significantly enriched in region-associated topics in fibroblasts (Fig. S3K), underscoring the potential regulatory roles of fibroblasts in the developing ChP.

### Homeostatic macrophage diversity within and across the developing ChP

The ChP is an entry point for immune cells and immune signaling into the central nervous system in health and disease^3,13^. Recent studies have identified a diversity of immune cells in the choroid plexus in adult^41^ and postnatal^42^ brain, yet the diversity of immune cells within and across the developing ChPs is not well understood. We identified eight subsets of immune cells: B cells, lymphocytes, macrophages, basophils, mast cells, dendritic cells, monocytes and neutrophils (Fig. 4A,B), each expressing specific immune regulatory genes encoding cytokines, chemokines, and complement components (Fig. S4A,B). For example, basophils expressed high levels of proinflammatory chemokines (*Ccl3, Ccl4, Ccl6, Ccl9*, Fig. S4A,B), suggesting that they may provide signals to trigger activation of signaling cascades necessary for leukocyte recruitment.

**Fig. 4.**
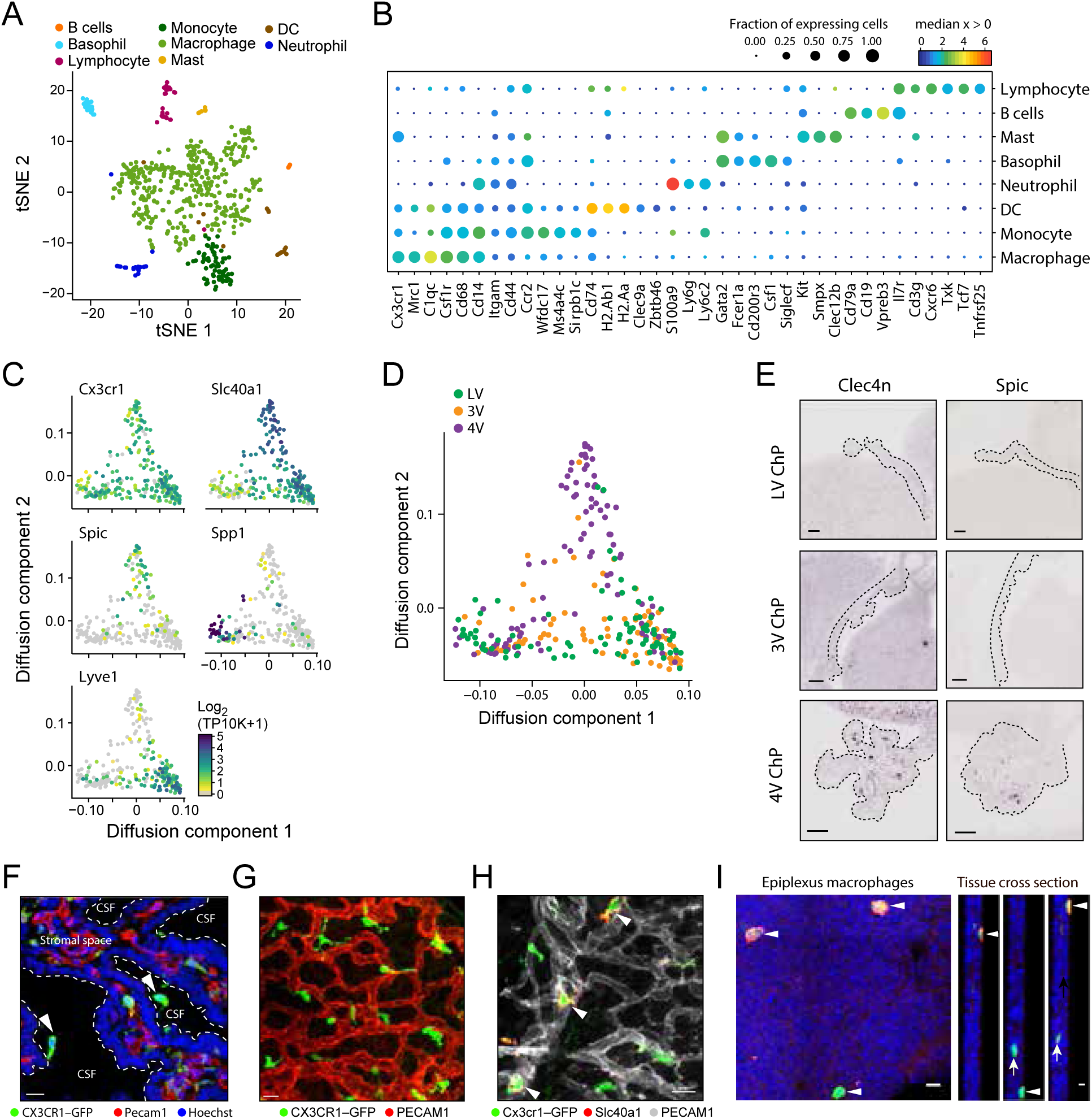
Immune cell diversity in the ChP and macrophage niches within and across ChPs. **(A)** Immune cell subsets in the ChP. tSNE of immune cell profiles, colored by cluster membership. **(B)** Marker genes for eight immune cell types. Median expression level in expressing cells (color) and proportion of expressing cells (circle size) of selected marker genes (rows) across immune cell subsets (columns). **(C)** Macrophages phenotypic heterogeneity across three archetypes. Diffusion map embedding of macrophages, colored by log_2_(TP10K+1) expression of a general marker (*Cx3cr1*) or each archetype specific gene (*Slc40a1, Spic, Spp1, Lyve1*). **(D)** Distinct macrophage archetype enriched in 4V ChP. Diffusion map embedding of macrophages (as in **C)** colored by ventricle. **(E)** Validation of macrophage subsets in 4V ChP. *In situ* hybridization (Genepaint) of *Spic* and *Clec4n* in LV, 3V, and 4V ChP (scale bar = 10 µm). Dotted line outlines the ChP. **(F)** *Cx3cr1^+^* macrophages in the sub-epithelial stromal space and supra-epithelial epiplexus positions in the ChP. Cortical section of LV from CX3CR1-GFP mouse co-stained with antibodies against endothelial marker PECAM1 (red) and nuclei (Hoechst, blue). Arrows: epiplexus cells. Scale=10 µm. **(G)** *Cx3cr1*+ macrophages in a tiled pattern closely associated with blood vessels in the ChP stromal space. Whole explants from CX3CR1-GFP mice co-stained with antibodies against PECAM1 (red). Scale=20 µm. **(H)** Diverse macrophages in the ChP stromal space. Whole explants from CX3CR1-GFP mice co-staining with antibodies against iron transporter FPN/SLC40A1 (red) and PECAM1 (grey). White arrow heads: macrophages localized to blood vessels that express the iron transporter. Scale=10 µm. **(I)** Diversity among epiplexus macrophages. Whole explants from CX3CR1-GFP mice co-stained with antibodies against hyaluronan receptor LYVE-1 (red). White arrow heads: LYVE1+ epiplexus macrophages in supra-epithelial positions (right panels: cross section. Scale bar = 10 µm in whole explants; 5 µm in cross-section images).

*Cx3cr1*^+^*Csf1R*^+^ macrophages were the largest class of immune cells in the ChP and showed graded gene expression patterns by diffusion map embedding (Fig. 4C, **Methods**), spanning states between three “archetypes”. All archetypes expressed *Mrc1* and *CD68*, which are involved in macrophage phagocytosis. Genes differentially expressed between archetypes included the hyaluronan receptor *Lyve1*^43^; *Spp1*, a potential regulator of phagocytosis in the brain^44^; *Slc40a1*, an iron transporter; and *Spic*, which marks red pulp macrophages^45^ and bone marrow macrophages (Kohyama et al., 2009) (Fig. 4C, S4C). A subset of *Slc40a1* macrophages expressed *Spic* and *Clec4n* (corresponding to recently described *Clec4n^+^* macrophages in postnatal ChP^42^), and was enriched in the 4V ChP (Fig. 4D,E, S4D). This highlights potential regionalization of ChP macrophages across the developing brain ventricles.

The archetypal expression patterns are also differentially associated with distinct spatial niches in the ChP, either basally under the epithelial cell monolayer or on the apical surface (*e.g.*, epiplexus positions: Fig. 4F). Imaging the distribution of macrophages using *Cx3cr1-GFP* transgenic mice (**Methods**) revealed a “tiled” pattern of cells in proximity with blood vessels (Fig. 4G). A subset of these macrophages stained for *Slc40a1/Ferroportin*, suggesting potential roles in regulating local iron homeostasis (Fig.s 4H). Of the *Cx3cr1^+^* macrophages located on the apical ChP surface, a subset expressed *Lyve1*, potentially revealing molecular identity of CSF-facing Kolmer/epiplexus cells (Fig.s 4I). Together, this immune cell diversity in the ChP provides a platform for investigating the multiple roles that immune cells have in the ChP in homeostasis and in disease conditions.

### Arterio-venous zonation and blood brain barrier protein expression in ChP vasculature

The choroidal artery delivers blood to each ChP^47^, but the identity, structure and development of vascular cell types in the ChP remain largely unknown. Using topic modeling, we found transcriptional programs associated with arterial (Topic 8), venous (Topic 11) and arteriolar gene expression (Topic 3) (Fig. 5A, S5A,B). In contrast to epithelial cells and fibroblasts, we found little evidence for regionalization of endothelial cells across ventricles (Fig. S5C,D).

**Fig. 5.**
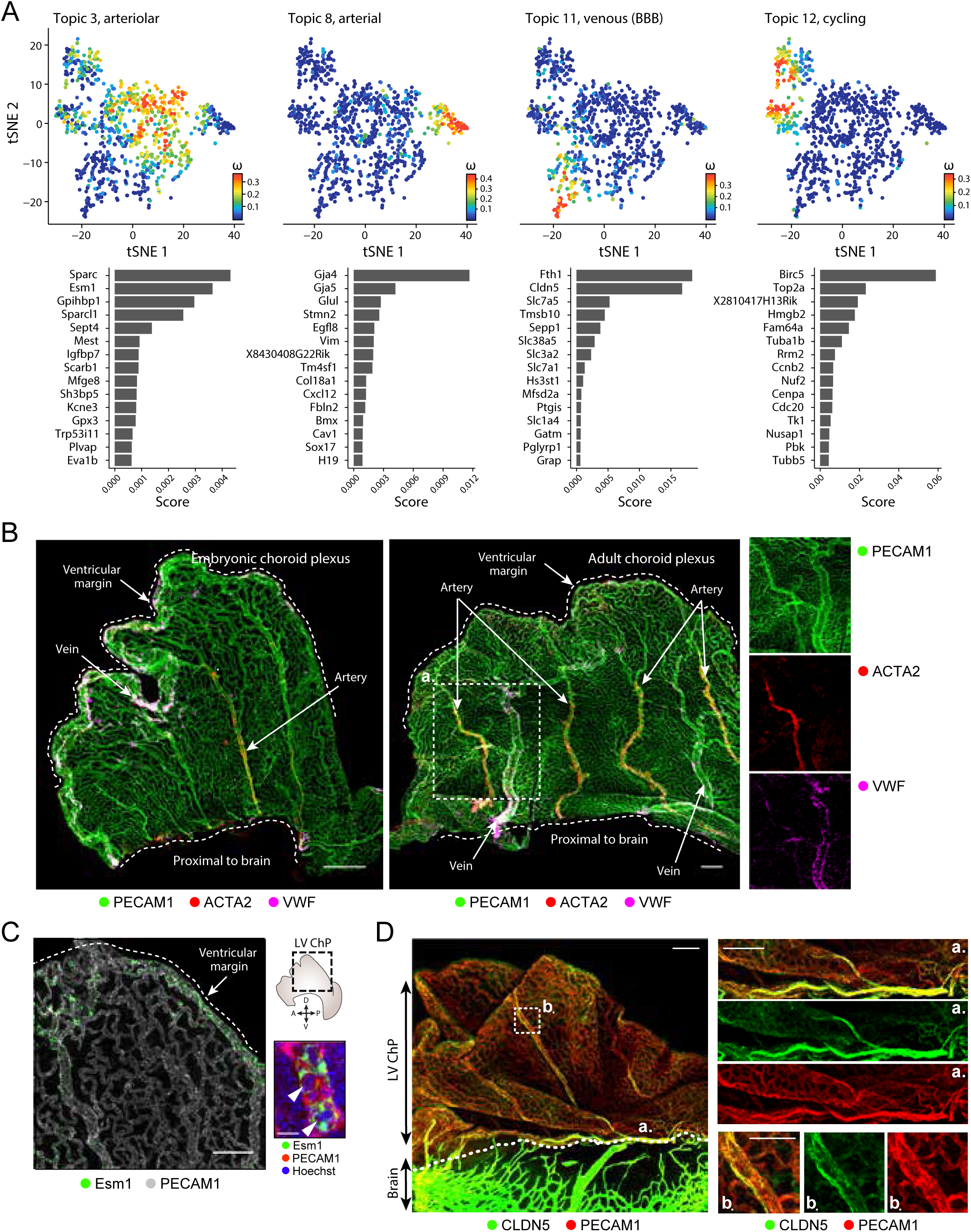
Vascular identity and BBB proteins are zonated within the ChP. **(A)** Endothelial cell transcriptional programs. For selected topics identified in endothelial cells, shown is tSNE of the cell profiles colored by topic’s weight per cell (top), and bar plot of topic scores for top ranked genes (bottom). Bold: Genes of interest. **(B)** Arterio-venous zonation in developing and adult ChP. LV ChP whole explants stained with antibodies against PECAM1 (general endothelial, green), ACTA2 (red) and VWF (magenta), marking the arterial (ACTA2+, VWF+) and venous (ACTA2-, VWF+) zones. Left: Emerging zonation in embryonic ChP. Middle: Clear arterio-venous organization in adult. Right: Insets from adult image. Scale bar = 100 µm. Zoomed inset scale bar = 50 µm. **(C)** Angiogenic zonation. Whole explant stained with antibodies against PECAM1 (grey) and the angiogenic marker ESM1 (green) enriched along the ventricular margin of the LV ChP. Scale = 100 µm. Top: Inset: Localization of the image within the explant and orientation within the brain. Arrows: anterior-posterior (A/P) and dorso-ventral (D/V) axes. Bottom: Co-staining for ESM1 (green), PECAM1 (red) and nuclei stain (Hoechst, blue). Scale=10 µm). **(D)** Blood brain barrier zonation. Whole explant stained with antibodies against CLDN5 (green) and PECAM1 (red) showing BBB marker expression at the root of the LV ChP, along vessels that run from the brain into ChP. Scale bar = 100 µm. Dotted line: barrier of the brain and the ChP. Right: Region a and b, scale bar = 20 µm.

We spatially mapped the developing arterio-venous zonation in the LV ChP in whole explants, by combining the pan-endothelial marker PECAM1 with markers for arteries (*Acta2+, Vwf+*) and veins (*Acta2-, Vwf+*)^48^, revealing arterial vessels oblique to the plexus and venous vessels along the ventricular margin of the tissue (Fig. 5B). By comparison, in the adult ChP there were regularly spaced arteries and veins across and expanded capillary-like network along the ventricular margin (Fig. 5B). This is reminiscent of the radially spreading vascular plexus in the developing retina^49^. The immature endothelial marker, *Esm1*, which marks angiogenic cells ^50,51^ was expressed in endothelial cells along the ventricular margin of the developing LV ChP (Fig. 5C), suggesting these cells may contribute to the expansion of LV ChP during maturation. Both *Esm1* and *Plvap*, which marks diaphrams of fenestrae in fenestrated endothelial cells^52^, scored highly in topic 3, suggesting the presence of developing fenestrations in the ChP as early as E16.5^53^ (Fig. 5A, S5E,F).

Surprisingly, the non-angiogenic endothelial cells expressed blood brain barrier proteins (topic 11), even though ChP lacks a classic blood brain barrier. These included *Cldn5* and *Mfsd2a*, which was expressed in cells with particularly high *Cldn5* expression^54^ (Fig. 5A, S5E). CLDN5 was expressed in endothelial cells *in situ* and was enriched in the brain and along the ChP region proximal to the brain (Fig. 5D). Such BBB-like expression may be a transient developmental feature of the ChP, analogous to that observed in the developing vasculature along retinal pigmented epithelium^55^.

### A network of potential cellular crosstalk in the ChP

Each of the diverse cell types in the ChP expressed a large number of genes encoding secreted proteins, many with potential to impact cell-cell interactions within the ChP or the composition of the CSF. In addition to epithelial cells, a recognized source of CSF-distributed factors, we identified additional potential cellular sources for proteins that have been measured in CSF (Fig. S6). For example, *Insulin like growth factor 2* (*Igf2*), which is secreted into the CSF and regulates progenitor proliferation in the developing cerebral cortex^6^, was highly expressed in endothelial and mesenchymal cell subsets in addition to epithelial cells (Fig. S7A). A cell-cell interaction network based on cognate receptor-ligand pairs (**Methods**), showed that endothelial, mesenchymal and immune cells potentially receive much greater signal input from other cell types compared to epithelial, neuronal and glial subsets. Further, mesenchymal cells expressed ligands for cognate receptors found on all the major cell types (Fig. S6B, S7B).

Examining cell specific interactions highlighted key potential roles for specific cell types. For example, basophils expressed colony stimulating factor 1 (*Csf1*), while macrophages and monocytes expressed its receptor *Csf1R*, suggesting a signaling axis for myeloid cell maturation (Fig. S6C), similar to basophil-macrophage communication in the lung^56^. In another example, basophils, and to a lesser extent mast cells, expressed *Il6*, whereas one of the IL6 receptors, *Il6st*, was predominantly expressed by mesenchymal cells, and another, *Il6ra*, was enriched in monocytes, macrophages and DCs (Fig. S6C). Basophils and mast cells also specifically expressed *Il1rl1* (ST2), the receptor for the alarmin *Il33*^57^, which in turn was expressed by fibroblasts and enriched in the 3V ChP (Fig. S6C). Finally, the growth factor receptors *Pdgfra* and *Pdgfrb* were uniquely expressed in fibroblasts and pericytes, respectively, while their ligands, *Pdgfa and Pdgfb*, were uniquely expressed by epithelial and endothelial cells, respectively (Fig. S6C). Together, these data highlighted a potential intercellular signaling niche within the ChP that may instruct overall ChP morphology and function.

### Maturation of the choroid plexus brain barrier in the adult brain

To assess to what extent the cellular diversity established within and across ChP of the developing brain is maintained in the adult, we performed single nucleus RNA-seq (snRNA-Seq)^58^ (**Methods**) of 52,629 nuclei across the three ventricles of the ChP from adult (4-6 month old) mice (n = 13 number of animals, in 3 pools for each ventricle, performed in two separate experiments) (Fig. S8A,B). We also profiled 9,845 nuclei from developing LV ChP from E16.5 embryonic brains (n=3 animals) for comparison. To relate adult and embryonic profiles we jointly clustered 62,474 nuclei from developing and adult ChP (**Methods**, Fig. S9B) and partitioned cells into ten clusters (Fig. S8B). As an alternative approach to compare adult and embryonic profiles, we used a random forest classifier (as in^58^, **Methods**, Fig. S9A). All cell types observed in our embryonic atlas were present in the adult ChP, along with additional adult-specific cell subsets (Fig. S8B, S9B-D), such as ependymocytes. While snRNA-Seq allowed us to overcome the challenge of dissociating adult tissue^58^, it under-represented immune cells (Fig. S8E, **Methods**).

Adult and embryonic nuclei showed differences in transcriptional profiles, proportions, maturation states and regionalization. For example, while in the embryonic brain we found glia and neuronal precursors and immature neurons (Fig. 1F), in the adult ChP we found distinct populations of mature neuronal cells (*Gria2* and *Kcnh7*, or the *Ddc* expressing subsets; Fig. S8C), and astrocyte-like cells expressing GFAP with ramified processes embedded in the choroid plexus stromal space (Fig. S9F). Other age-specific differences included lack of proliferating and ciliogenesis epithelial cells in the adult cell types, suggesting low proliferation under homeostatic conditions in the adult ChP (Fig. S8D). Finally, mesenchymal cells clustered into three subtypes, an embryonically enriched cluster (mesenchymal 1), an adult enriched cluster (mesenchymal 2), and one shared cluster (mesenchymal 3) (Fig. S8B, S9B).

Interestingly, we found regionalized expression in adult epithelial cells, as also observed during development (Fig. S9G). Some genes maintained regionalized expression as in the embryo (e.g. *Wls, Tbc1d1, Sulf1*, and *Ins2*, Fig. S8E, S9H), such as *Ins2*, which retained its enriched expression in the 3V ChP epithelial cells (confirmed using single molecule fluorescence *in situ* hybridization (smFISH), Fig. S8E). Other genes lost expression altogether in adult epithelial cells, including *Penk* and *Shh*, which were no longer detected in the adult ChP (Fig. S9H)^4^. Finally, newly regionalized genes emerged in the adult epithelial ChP, including *Ttr*^4^ and *Slc35f1* (Fig. S9H). There was also some inconclusive ventricle specific regionalization of mesenchymal cell programs (Fig. S9I). Overall, there were adult-specific cellular subtypes and regionalized gene expression programs, along with retention of embryonic properties of the ChP, which altogether, reflect conserved and maturing functions of the ChP-CSF system.

## DISCUSSION

The choroid plexus (ChP) is a critical yet understudied brain barrier. The lack of a cellular and molecular parts list for the ChP has been a major obstacle to dissecting its roles in instructing brain development and health and unlocking its therapeutic potential. Our comprehensive cell atlas allows us to chart all major cell classes in the developing and adult ChP, including epithelial, endothelial, mesenchymal (mural and fibroblasts), immune (innate and adaptive), neuronal and glial cells. We characterized similarities and distinctions across brain ventricles and developmental states. Our extensive analyses and validations provide a molecular launchpad for defining the functional roles of each cell type and their interactions.

Establishing this resource required new solutions to several technical challenges unique to the choroid plexus, including tissue isolation, dissociation, and imaging. While gene expression has been analyzed previously from the LV and/or 4V ChP^4,7,22^, such analyses were not performed for the 3V ChP, likely due to the difficulty of reliably dissecting this tissue. Here, we optimized micro-dissection techniques to isolate ChP from each ventricle, with minimal contamination from adjacent brain tissue. Next, we optimized enzymatic dissociation of embryonic tissue into single cells by employing combinations of enzymes, temperatures, and mechanical dissociation. We isolated nuclei from adult tissue, which failed to yield viable cells by dissociation. Analyzing embryonic tissue at both the single cell and single nucleus level allowed us to assess some of the similarities and differences between embryonic and adult ChP tissues.

Our work builds on previous findings^4^ to demonstrate that epithelial expression programs in the developing and adult brain are ventricle-specific. In the adult 4V ChP, cells retain differential expression of classic patterning genes (*e.g.*, *Hoxa2, Hoxb3os*, *Meis1*) as transcriptional memories of their early hindbrain development along the rostro-caudal body axis. Such regionalization may help drive ventricle-specific synthesis of factors to be secreted into the CSF on extracellular vesicles for delivery to distal targets^59,60^. Alternatively, local paracrine signaling networks may be supported by *Shh* receptor *Ptch1* enrichment in 4V ChP stromal cells and epithelial cells, in addition to earlier reports that showed that *Shh-Ptch1* pericyte signaling instructs the coordinated vascular outgrowth of the fourth ventricle ChP^61^.

Our data revealed several putative signaling axes across cell types, highlighting endothelial and mesenchymal cells as additional potential sources of signals within the ChP. These results open new hypotheses about paracrine signaling within the ChP that may contribute to functions beyond immune cell recruitment and tissue development. Our cell atlas provides a first opportunity to identify specific cell types within and across ventricles responsible for the production of secreted factors that promote the health and growth of the brain^1,3,62^. Previous studies predicted ChP epithelial cells to be the primary producers of a multitude of secreted factors for distribution throughout the developing brain via the CSF. However, we show that essentially all major ChP cell types, and in particular ChP fibroblasts, also express many secreted factors. It remains to be determined if these secreted factors of non-epithelial origin are restricted to targets within ChP stromal space, or whether transport mechanisms allow them to be delivered into the CSF.

In particular, we found that *Ins2* expression was enriched in 3V ChP epithelial cells, suggesting the ChP as a potential internal source of insulin for the brain. While CSF-insulin levels are well documented^63^, the potential for a brain-derived source of functional insulin has been long debated^64^. Receptors and signaling machinery for insulin are found widely throughout the brain, the hypothalamus and ventral third ventricle – major centers involved in regulation of body metabolism – are located in the immediate neighborhood of the 3V ChP and could be nearby target sites^65,66^, where local fluid flows distribute factors^67^. ChP-derived insulin could also act more distally, including on neural progenitor cells to regulate neurogenesis (as has been shown for IGF family members ^6^).

Ventricle-specific regionalization of mesenchymal cells in developing brains is accompanied by ventricle-specific expression of different ECM components including collagens, and could contribute to differential tissue stiffness and elasticity, which is apparent upon tissue dissection. For other tissues of the body such as the developing muscle and bone, ECM properties impart instructive cues to stem cells^68^, suggesting that spatially heterogeneous ECM niche in ChP may guide cell-cell interactions and even cell migration^69^.

Our profiles of embryonic ChP immune cell populations under homeostatic conditions are largely in agreement with recent single cell profiling studies of adult ChP macrophages, and revealed spatial organization of several macrophage archetypes across ventricles and within the choroid plexus. The vast majority of our ChP macrophages shared 5 of 6 signature genes (*Mrc1, Ms4a7, Pf4, Stab1, Fcrls*) recently described for adult ChP macrophages^41^ and are *Sall1* negative^44^. Our analyses further revealed unique spatial positioning for *Spic*-expressing cells in the 4V ChP, where they localized to the medial core of the ChP. *Spic* expression correlated with that of *Clec4n*, a proposed marker of juvenile ChP macrophages^42^. *Slc40a1/Fpn-*expressing macrophages closely positioned along larger blood vessels. In other epithelia such as the intestine, *Slc40a1/Fpn* is an essential iron exporter that removes iron from cells into the blood^70^. Our findings suggest that *Slc40a1/Fpn-*expressing macrophages may participate in maintenance of brain iron homeostasis essential for brain development^71^.

Overall ventricle-specific regionalization of epithelial cells was maintained in the adult tissue, many of the genes underlying regionalization differed from those in embryonic samples. While *Ins2* retained in 3V ChP epithelial cells of the adult, some embryonically regionalized genes were no longer regionalized in the adult brain, and vice versa. Except for proliferating and differentiating epithelial cell states in the developing embryo, our conservative clustering did not identify distinct clusters of cells within each ventricle of either embryonic or adult, but rather, uncovered ventricle-specific expression, as reflected by “topics” enrichment. Topics as well as graded gene expression patterns, particularly across 4V epithelial cells, may arise from the segmental development of the 4V ChP from distinct rhombomeres^20,72^. Further heterogeneity in all cell types may emerge in response to state-dependent changes including circadian rhythms^10^ or in disease^41,44^.

Our study provides a first comprehensive map of the molecular, cellular and spatial diversity of the ChP from each ventricle of the developing and adult brain. Revealing a rich network of cells, with expression regionalization across ventricles in epithelial and mesenchymal cells, and suggesting a previously unappreciated role for mesenchymal, vascular and immune cells in shaping the tissue and potentially the CSF. These data will facilitate the design of tools to access and control specific cell populations in the ChP by genetic, optogenetic, and chemogenetic means. Since the ChP-CSF system is implicated in a growing number of neurologic conditions, our dataset offers molecular insight that will accelerate future studies investigating the lifelong, active regulation of the ChP brain-body barrier in health and disease.

## SUPPLEMENTARY FIG. LEGENDS

**Fig. S1.**
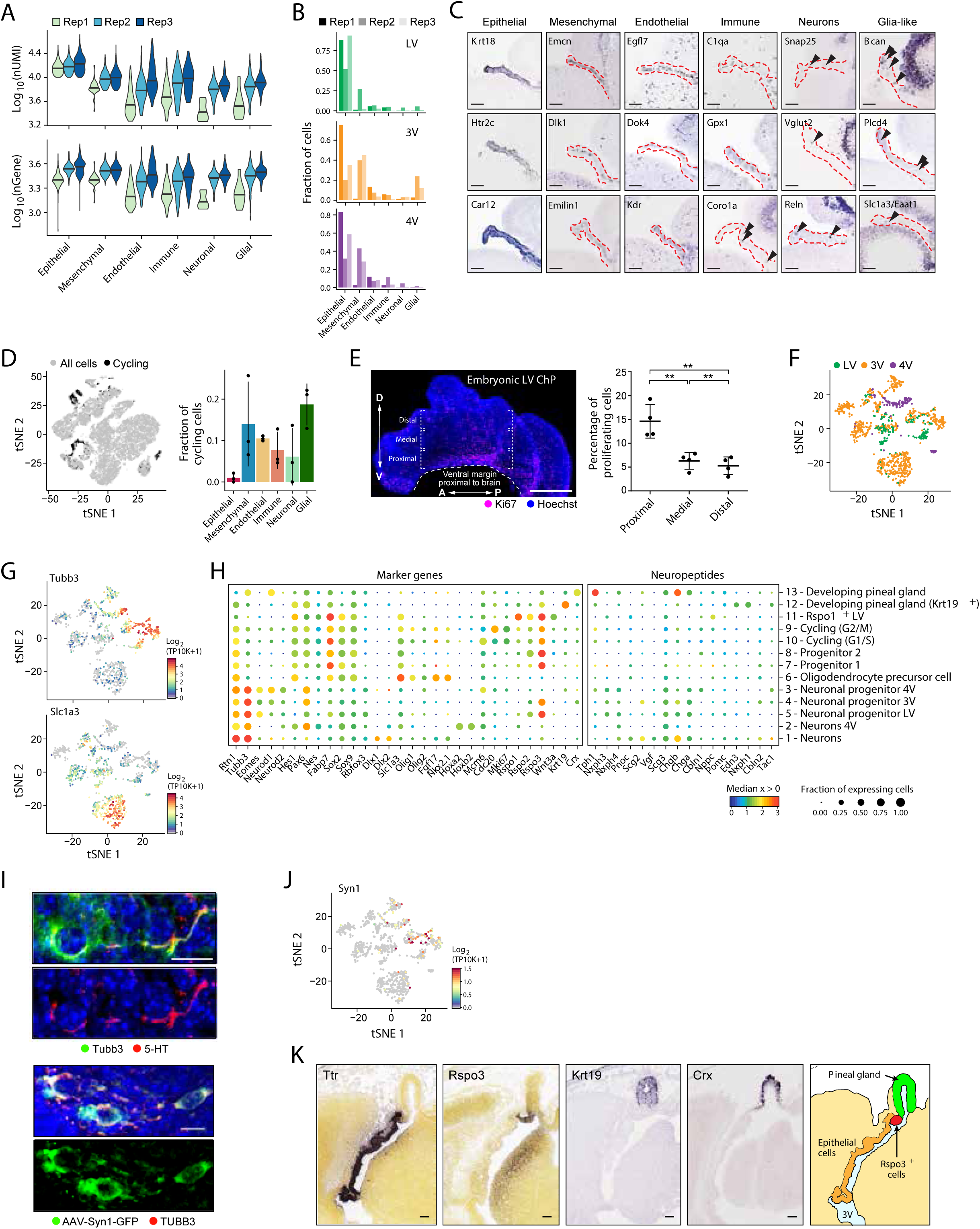
Cellular markers, identity and organization within the ChP. **(A)** Cell quality of scRNA-seq. Distribution of top: log_10_(nUMI) and bottom: log_10_(nGene) (y axis) across cell types, grouped and colored by replicate (x axis). **(B)** Proportions of cells captured from each ChP. Fractions (y axis) of each major cell type identified (x axis) per ventricle and experiment. **(C)** Canonical markers identify cellular localization in LV ChP. *In situ* hybridization images from the LV ChP from Genepaint of cell-specific marker genes. (Scale bar = 100µm). **(D,E)** Proliferating cell niche proximal to the brain. (**D**) Proliferating cells. Left: tSNE (as in Fig. 1C) of cell profiles (dots), with cells classified as cycling colored black (**Methods**). Right: Mean fraction of cycling cells (y axis) in each cell type (x axis), and in each replicate (dots). Line: standard deviation (SD). **(E)** Proliferating cell niche. Left: KI67 staining (magenta) of whole explant of LV ChP. Double headed arrows: A/P and D/V axes. Right: Percent of proliferating cells (KI67^+^, y axis) in proximal, medial and distal regions of the LV ChP (x axis, relative to the ventral margin of the ChP near the brain) (n=4, Scale bar = 500 µm; one-way ANOVA, ** p<0.01. Error bars indicate S.E.M. Scale bar = 500 µm). **(F)** tSNE of neuronal and glia-like cell profiles (dots, as in Fig. 1F), colored by ventricle. **(G)** tSNE of neuronal and glia-like cell profiles (dots, as in Fig. 1F), colored by log_2_(TP10K+1) expression of top: *Tubb3* (neuronal marker) and bottom: *Slc1a3* (glial marker). **(H)** Neuronal/glia-like subset markers and neuropeptides. Median expression level in expressing cells (color) and proportion of expressing cells (circle size) of selected genes (columns) in neuronal and glia-like subsets (rows). **(I)** Molecular expression in LV ChP neurons. Top: Representative image of a neuron stained with serotonin anti-sera(5-HT) (red) and anti-TUBB3 (green) antibody. Bottom: Representative image of neurons targeted by AAV expression of GFP driven by promoter of neuronal marker gene, *Syn1*, counterstained with anti-TUBB3 antibody. Across images: nuclei stained (blue). Scale bar = 10 µm. **(J)** Neuronal cells in the LV ChP. tSNE of neuronal and glia-like cell profiles (dots, as in Fig. 1F), colored by log_2_(TP10K+1) expression of *Syn1*. **(K)** Organization of the developing 3V ChP. *In situ* hybridization images (Genepaint) of marker genes for developing pineal gland clusters (*Krt19, Crx*), of progenitor cells (*Rspo3*) and of epithelial cells (*Ttr*), localizing the progenitors adjacent to the pineal gland.

**Fig. S2.**
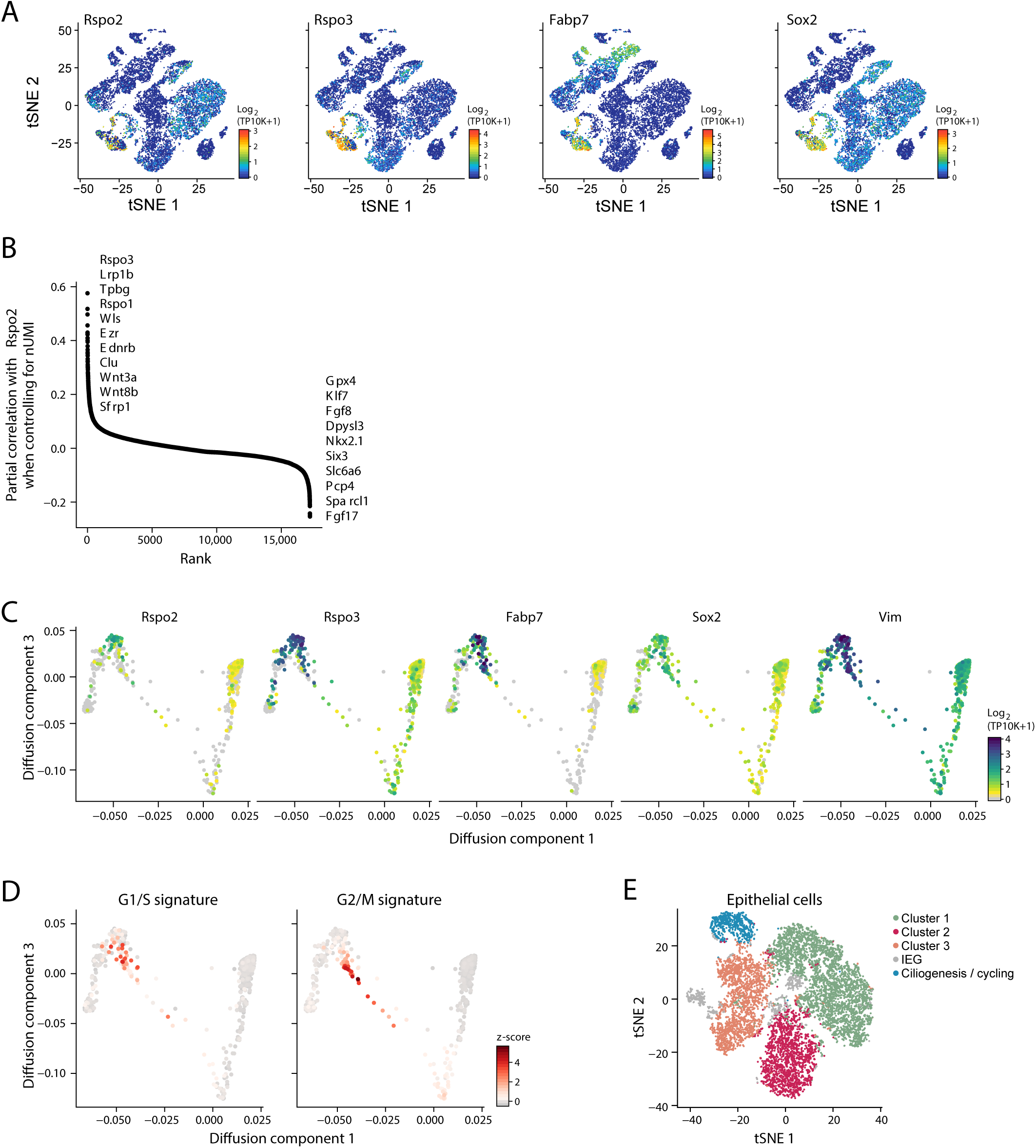
Common progenitors of neural epithelium divide to become mature epithelial cells. **(A-C)** Stem-like marker gene expression. **(A)** tSNE of all cell profiles (dots, as in Fig. 1C), colored by log_2_(TP10K+1) expression of denoted genes. **(B)** Partial gene expression correlation with *Rspo2* expression (y axis) by rank (x axis), across cells used for diffusion map (as in Fig. 2A). **(C)** Diffusion map (as in Fig. 2A), colored by log_2_(TP10K+1) expression of denoted genes. **(D)** Proliferating cells on the inferred trajectory. Diffusion map (as in Fig. 2A), colored by signature scores of left: G1/S and right: G2/M gene sets. **(E)** Epithelial cell clusters. tSNE of epithelial cell profiles (dots), colored by cluster membership.

**Fig. S3.**
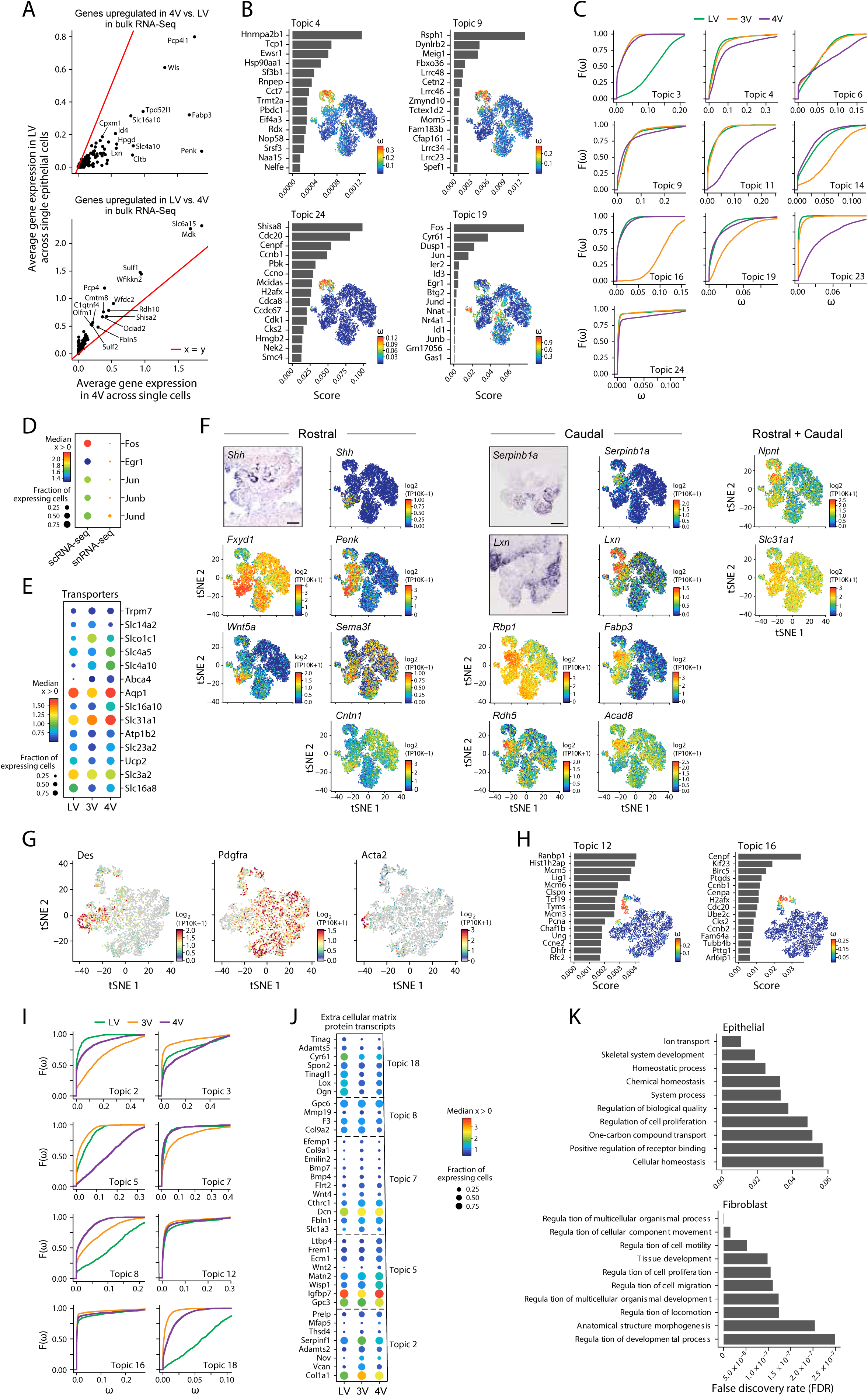
Regionalized transcriptional programs in epithelial and mesenchymal cell populations. **(A)** Single-cell data agrees with previous bulk RNA-seq data. Differentially expressed genes between LV and 4V in bulk RNAseq^4^ (Top: up-regulated. Bottom: down regulated) averaged across single cell in 4V ChP (x axis) and LV ChP (y axis). Red line indicates x=y. All genes show a coordinated difference between the datasets. **(B)** Transcriptional programs associated with immediate early gene expression (Topic 19, bottom right) and ciliogenesis (Topic 24, bottom left) and cell cycle (Topic 4, 9, top) in epithelial cells. For each topic, shown is a bar plot of topic scores for top ranked genes (left), and tSNE of the cell profiles (as in Fig. 3A) colored by topic’s weight per cell (right). **(C)** Ventricle associated transcriptional programs in epithelial cells. Empirical cumulative distribution (y axis) of topic’s weight (x axis), grouped and colored by ventricle. **(D)** Immediate early gene (IEG) expression in scRNA-seq and snRNA-seq data. Median expression level in expressing cells (color) and proportion of expressing cells (circle size) of selected genes (columns) in our scRNA-seq and snRNA-seq data (rows). **(E)** Regionalized expression of transporters in epithelial cells. Median expression level in expressing cells (color) and proportion of expressing cells (circle size) of transporters within top 50 features (rows) across ventricles (columns). **(F)** Rostral and caudal gene expression pattern in 4V ChP epithelium. tSNE of epithelial cell profiles (dots), colored by log_2_(TP10K+1) expression of genes shown in Fig. 3E, F. **(G)** Mesenchymal cells partition into fibroblast and mural cells (pericytes and smooth muscle cells). tSNE of mesenchymal cell profiles (dots), colored by log_2_(TP10K+1) expression of marker genes for: pericytes (*Des*), smooth muscle actin positive cells (*Acta2*) and fibroblasts (*Pdgfra*). **(H)** Transcriptional programs associated with cell cyle (Topic 12,16) in mesenchymal cells. For each topic, shown is a bar plot of topic scores for top ranked genes (left), and tSNE of the cell profiles (as in Fig. 3A) colored by topic’s weight per cell (right). **(I)** Ventricle associated transcriptional programs in mesenchymal cells. Empirical cumulative distribution (y axis) of topic’s weight (x axis), grouped and colored by ventricle. **(J)** Regionalized expression of extracellular matrix (ECM) protein in mesenchymal cells. Median expression level in expressing cells (color) and proportion of expressing cells (circle size) of ECM genes within top 50 features (rows) across ventricles (columns). **(K)** Functional enrichment in regionalized transcriptional programs in epithelial and mesenchymal cells. GO analysis enrichment of union of top 50 features of regionalized topics are shown for epithelial cells (top) and mesenchymal cells (bottom). FDR (x axis) of top ten significant biological processes (y axis) are shown.

**Fig. S4.**
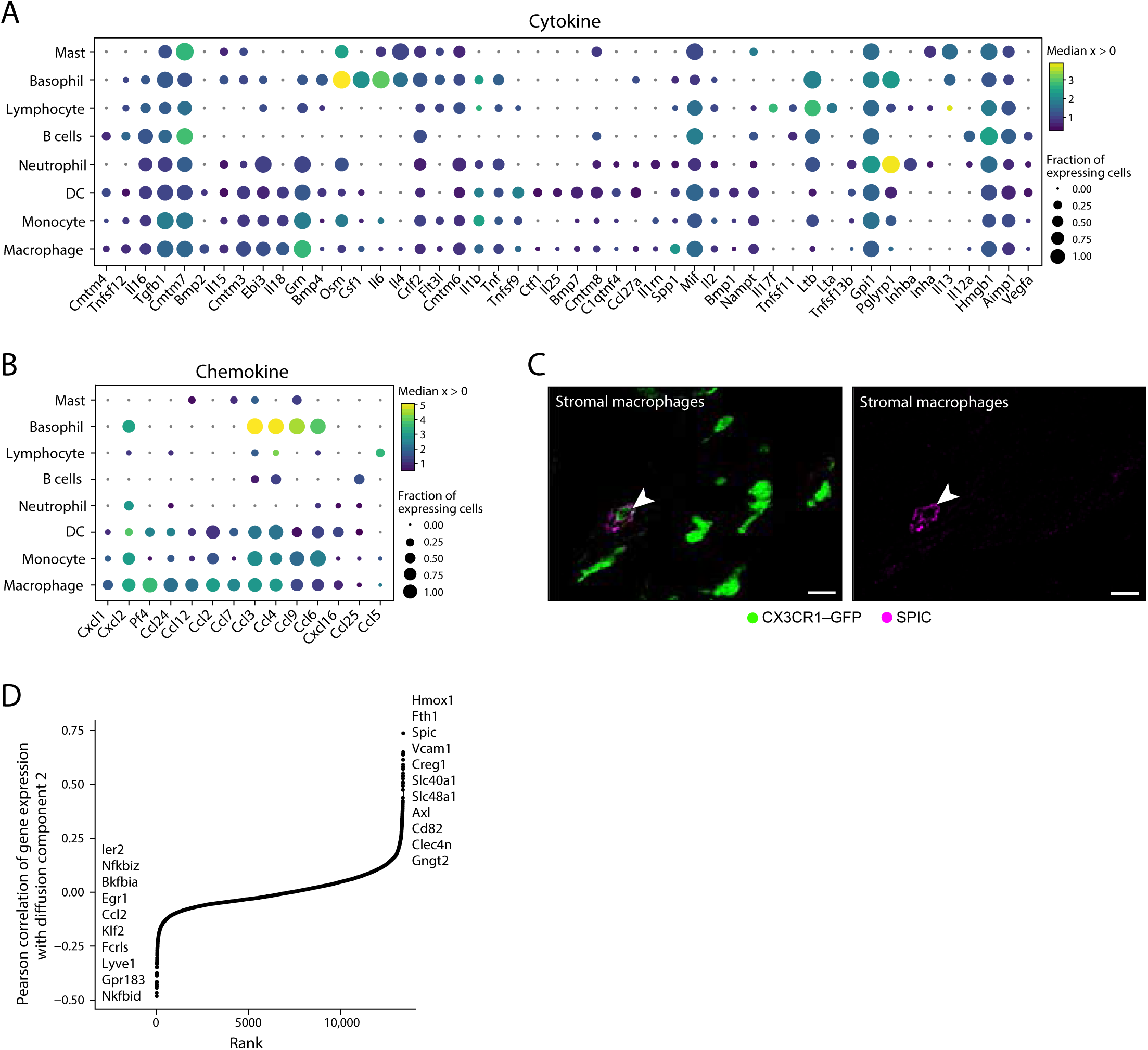
Cytokine and chemokine expression across ChP immune cells. **(A-B)** Gene expression of cytokines and chemokines across immune cell subsets. Median expression level in expressing cells (color) and proportion of expressing cells (circle size) of cytokines **(A**, columns) and chemokines **(B**, columns) in immune cell subsets (rows). **(C)** SPIC^+^ macrophages in ChP stromal space. LV ChP explants from CX3CR1-GFP mice stained with SPIC antibody (magenta). White arrow heads: SPIC^+^ macrophages. Scale bar = 20µm. **(D)** Correlated gene expression in *Spic* expressing macrophages. Partial gene expression correlation (normalized for nUMI, y axis) with cell scores for diffusion component 2 from diffusion map (as in Fig. 4C).

**Fig. S5.**
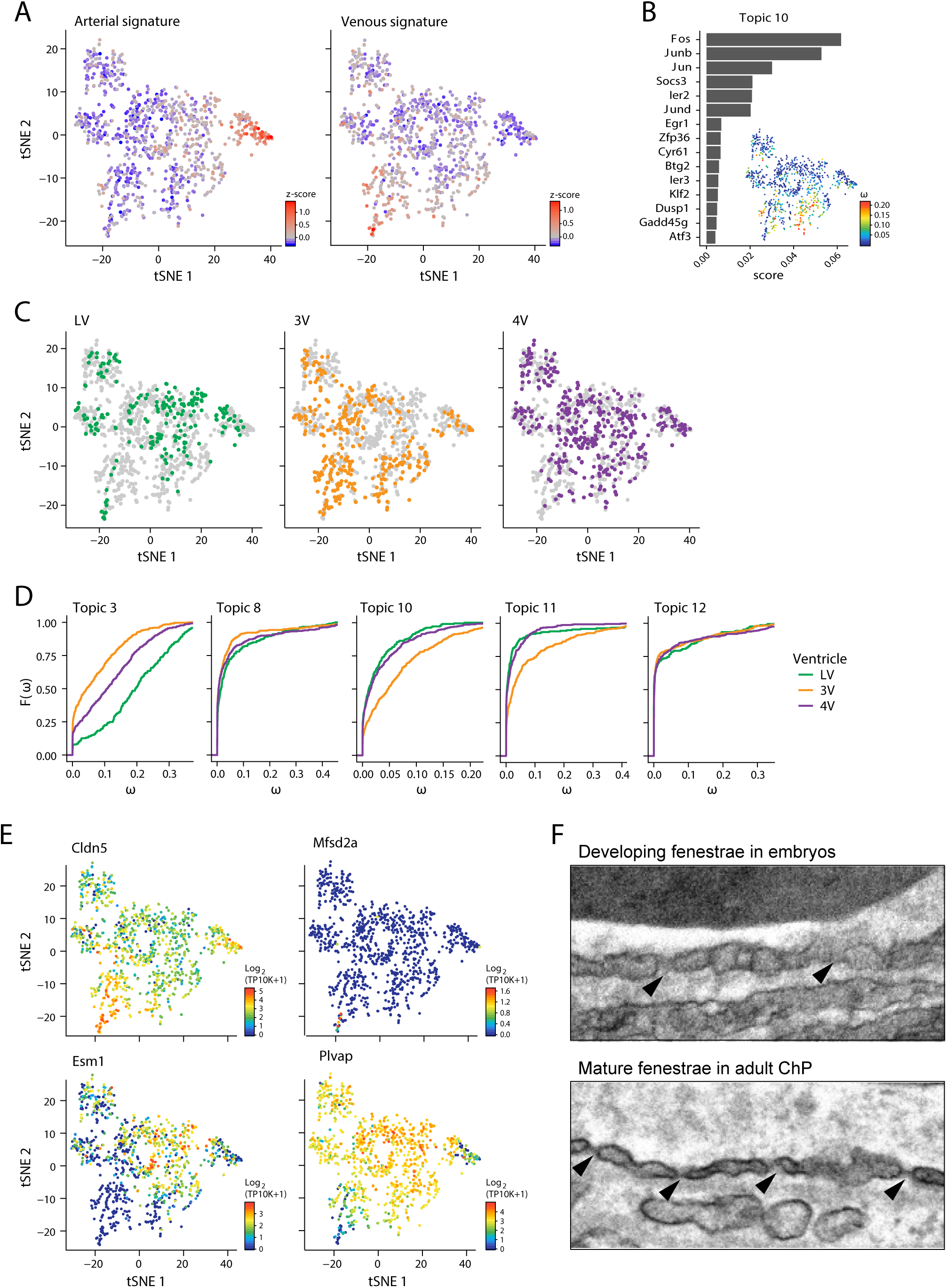
Molecular identities of ChP endothelial cells. **(A)** Arterial and venous transcriptional programs in endothelial cells (EC). tSNE (as in Fig. 5A) of EC profiles (dots), colored by signature scores for arterial (left) and venous (right) gene sets (signatures taken from^48^). **(B)** Transcriptional program associated with immediate early gene expression. Shown is a bar plot of topic scores for top ranked genes (left), and tSNE of the cell profiles (as in **A**) colored by topic’s weight per cell (right). **(C)** tSNE (as in **A**) of cell profiles (dots), colored and faceted by ventricle. All EC are shown in grey as background. **(D)** Empirical cumulative distribution (y axis) of topic’s weight (x axis), grouped and colored by the ventricle. **(E)** Blood brain barrier gene expression in EC. tSNE (as in **A**) of endothelial cell profiles (dots), colored by log_2_(TP10K+1) expression of BBB genes: *Cldn5* (top left), *Mfsd2a* (top right), *Esm1* (bottom left) and *Plvap* (bottom right). **(F)** Transmission electron microscopy image of developing fenestrae in the embryonic vessels (top, black arrow heads), as compared to well-defined fenestrae in the adult endothelia (bottom, black arrow heads). Scale bar = 100 nm.

**Fig. S6.**
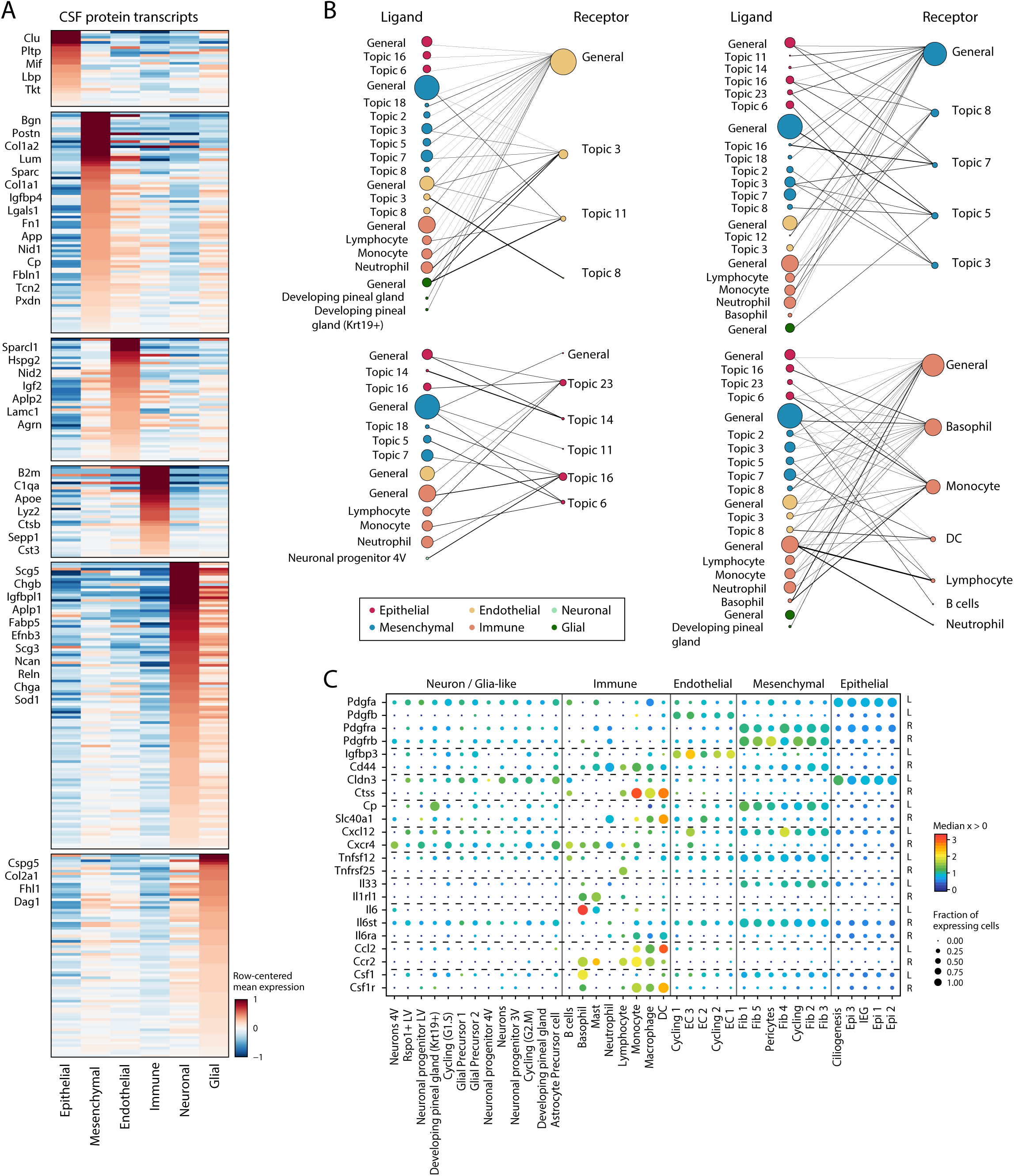
Mesenchymal, endothelial and immune cells contribute to cellular cross-talk in ChP. **(A)** Many ChP cell types express genes encoding secreted factors. Mean expression (color bar, row centered) of genes (rows) coding for proteins measured in CSF from embryonic mice^4^ in each of six major cell types (columns). Genes are sorted by expression level in the cell type in which they were maximally expressed. Genes of interest are marked on the side. **(B,C)** Potential roles for mesenchymal, endothelial, and immune cells in cell-cell interactions. **(B)** Bipartite graph of cellular network of ligand (left) and cognate receptor (right) pairs. Nodes: sets of ligands (left) or receptor (right) genes, which are either cell type specific or subset specific (identified by differential expression for immune cells and neuronal /glia-like cells, or by top 50 scoring genes of topics, **Methods**). Node color: cell type; Node size: degree in the full network. **(C)** Expression of examples (from B) of key ligand-receptor pairs (rows; groups of receptors and cognate ligands are separated by vertical dashed lines) across cell subtypes (columns). Dot size: fraction of expressing cells; color: median expression in expressing cells.

**Fig. S7.**
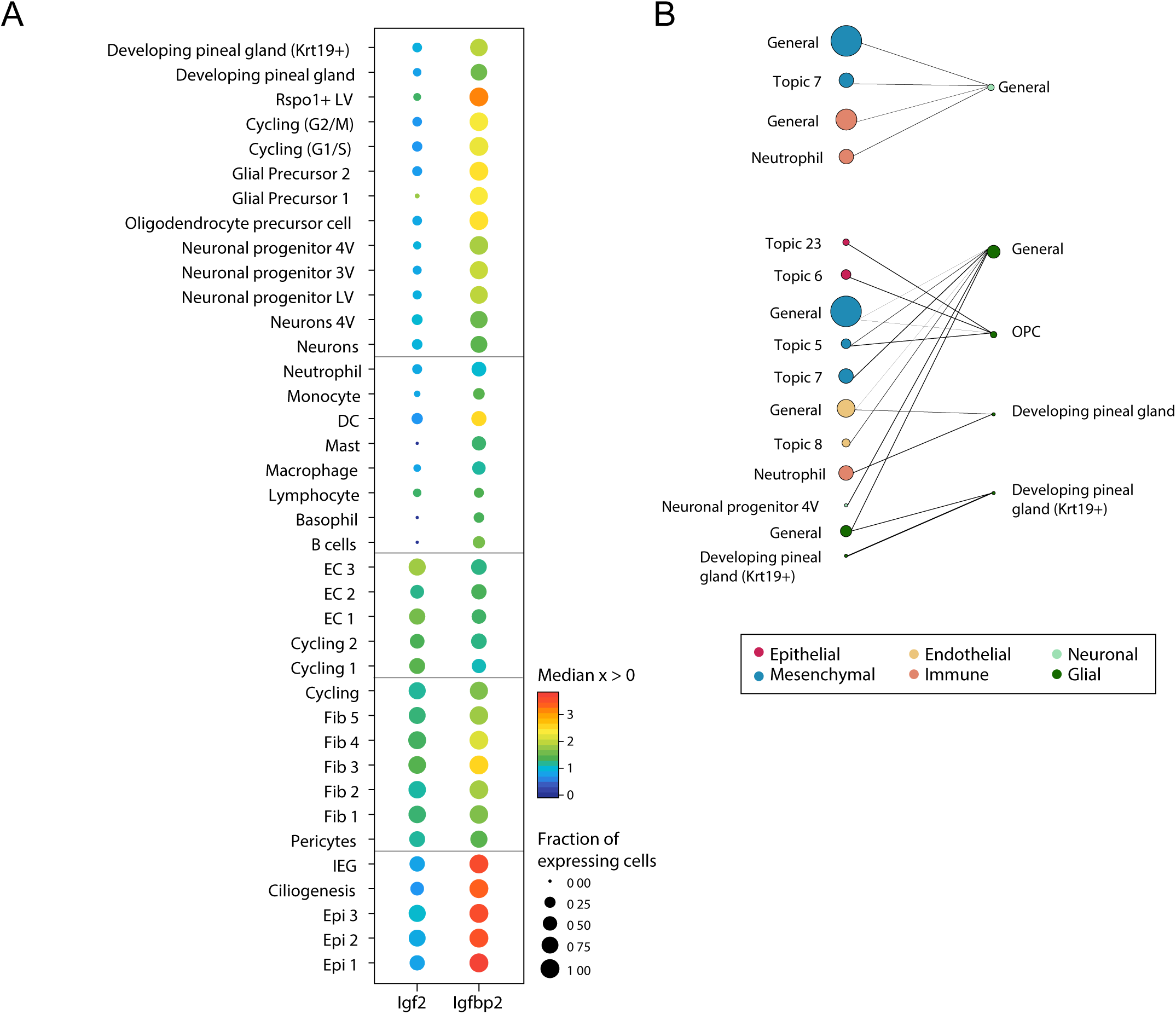
Neuronal and glia-like cells do not play significant contribution in cell-cell crosstalk in the developing ChP. **(A)** Expression of example secreted genes across all cells. Median expression level in expressing cells (color) and proportion of expressing cells (circle size) of denoted genes (columns) in all cell type subsets (rows). **(B)** Bipartite graph of cellular network for neuronal (top) and glial cells (bottom).

**Fig. S8.**
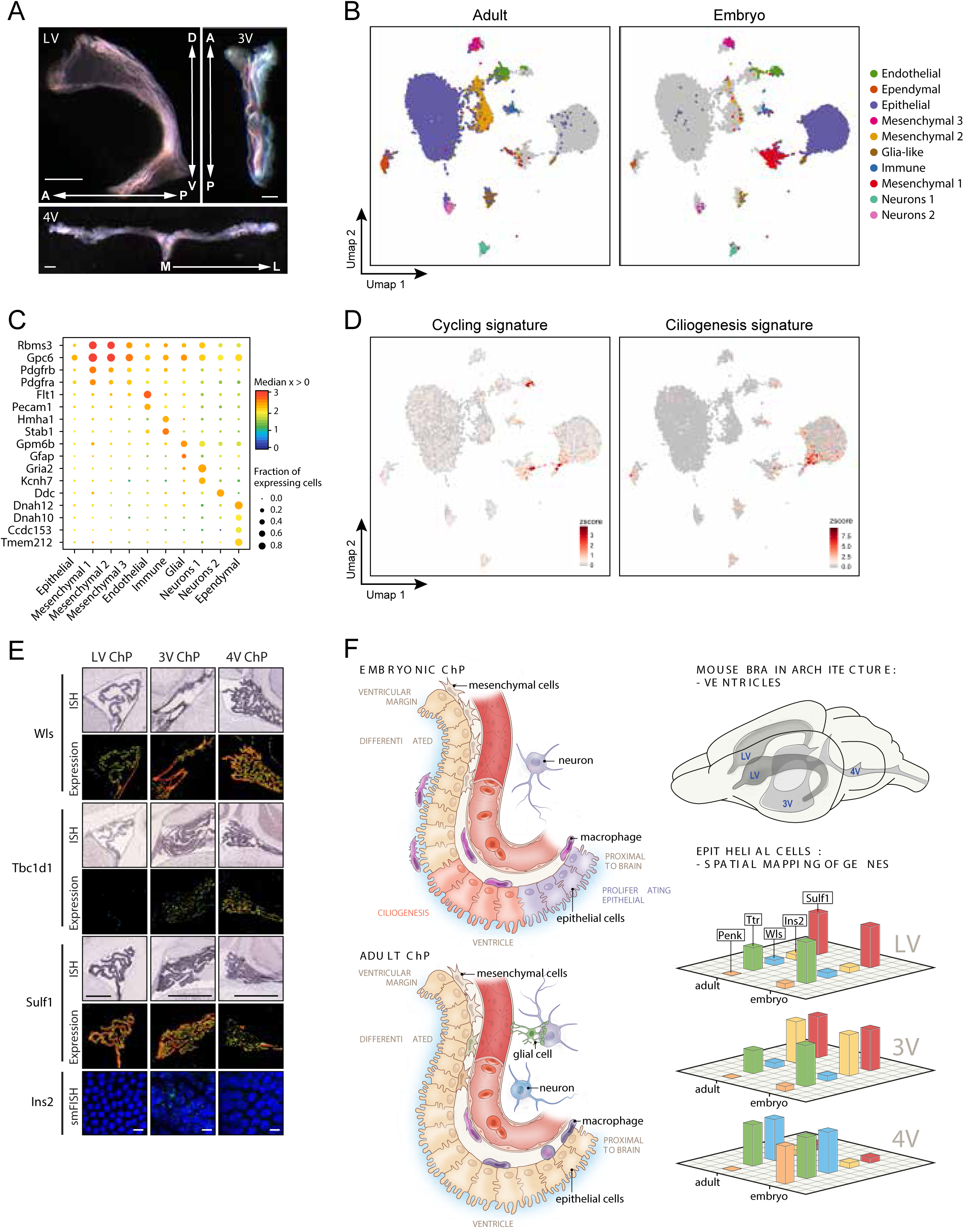
Maturation of ChP brain barrier in adulthood. **(A)** Adult ChP. Whole explant images of the adult ChPs (Scale bar = 1 cm). Arrows: anterior-posterior (A/P), dorso-ventral (D/V) and medial-lateral (M/L) axes. **(B)** snRNA-Seq from embryo and adult ChP. UMAP embedding of 29,727 sampled single nucleus profiles (**Methods**, dots, adult: n=13 mice, processed in 3 pools of per ventricle, embryo: n=3 mice, processed in 1 pool of LV ChP), colored by annotated cell type (epithelial cell clusters were merged *post-hoc*), in either adult (left) or embryo (right). **(C)** Cell type marker genes in adult. Mean expression in expressing cells (color) and proportion of expressing cells (dot size) of selected genes (row) across the major cell populations (column). **(D)** Cell proliferation and ciliogenesis programs are specific to embryonic ChP cells. UMAP embedding (as in **B**) with cells colored by signature score of cell cycle (left) and ciliogenesis (right). **(E)** Regionalized expression of adult epithelial cells across ventricles conserved from the embryo. Regionalized gene expression in epithelial cells by ISH images (top) from Allen Brain Atlas^73^ or smFISH (bottom), across the adult LV ChP, 3V ChP and 4V ChP (columns). **(F)** Model of age-dependent changes in ChP cell types and tissue organization (left), and ventricle-specific gene expression in epithelial cells (right).

**Fig. S9.**
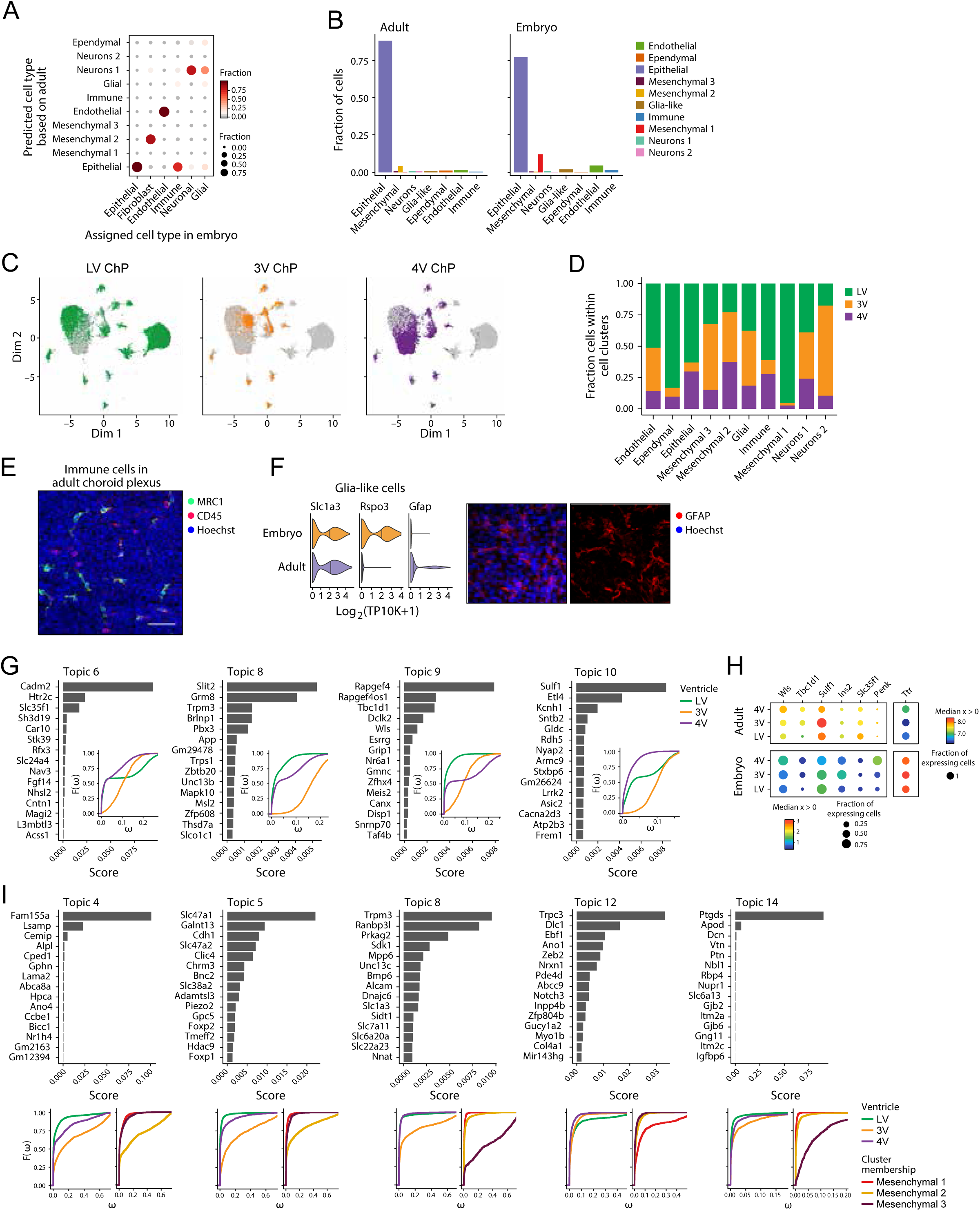
Molecular heterogeneity across adult ChP cell types. **(A)** A random forest classifier assigns cell types across age. Fraction of cells (color and circle size) per cell type in embryo (column) that were mapped to adult clusters (row) (**Methods**). Columns add up to 1. **(B)** Proportions of cells captured for embryo and adult in snRNA-seq data. Fractions (y axis) of each major cell type identified (x axis, color) across age. **(C)** UMAP embedding (as in Fig. S8B) of cell profiles (dots), colored and faceted by ventricle. All cells shown in grey as orientation. Adult ChP was sampled form all three ventricles, and embryo from LV only. **(D)** Composition of each cell cluster (x axis) by ventricle (color). **(E)** Immune cells in the adult ChP. Whole explant imaging of the LV ChP stained with antibodies against MRC1 (green) and CD45 (red). Nuclei in blue. Scale bar = 50 µm. **(F)** GFAP^+^ cells identified in the adult ChP. Left: gene expression of glial cell markers (columns) in embryonic and adult glia-like cells (rows). Right: Immunohistochemistry with antibody against GFAP (red, nuclei in blue) in the adult LV ChP explants (Scale bar = 10 µm). **(G)** Regionalized transcriptional programs in adult epithelial cells. For each topic that is differentially weighted between ventricles, shown is a bar plot of topic scores for top ranked genes (left), and empirical cumulative distribution (right) of topic’s weight, grouped and colored by ventricle. **(H)** Regionalized gene expression in epithelial cells across age, some examples. Median expression level in expressing cells (color) and proportion of expressing cells (circle size) of denoted genes (columns) across ventricle and age (rows). **(I)** Transcriptional programs in adult mesenchymal cells. For selected topics, shown is a bar plot of topic scores for top ranked genes (top), and empirical cumulative distribution (bottom) of topic’s weight, grouped and colored by ventricle (left) and mesenchymal cluster (right).

## MATERIALS AND METHODS

### Animal models

All mouse work was performed accordance with the Institutional Animal Care and Use Committees (IACUC) and relevant guidelines at the Boston Children’s Hospital, Broad Institute and MIT, with protocols 17-10-3547R.

Embryonic day (E) 16.5 embryos were obtained from time-pregnant CD-1 dams for all embryonic ChP analyses. Adult ChP were harvested from 4-6 months old CD-1 males. All animals were housed under 12hr/12hr day night cycle with access to standard chow and water ad libitum. Embryonic day (E) 16.5 Cx3cr1-GFP mice were used to visualize macrophage distribution in the embryonic ChP.

CD1(ICR)– Charles River Labs Strain 022.

Cx3cr1-GFP (catalog #: 005582, The Jackson Laboratory)

### Embryonic ChP tissue dissection

Whole ChP from each ventricle were harvested using #5 forceps and fine-dissection scissors. To collect the 4V ChP, the developing hindbrain was separated from the mid- and forebrain structures using a scalpel. Next, using a scalpel the cisterna magna was exposed by gently pushing away the developing cerebellum to expose the 4V ChP. Lateral arms and medial core of the 4V ChP were teased away using forceps. Next, the 3V and LV ChP were collected from the rest of the developing brain structures. 3V ChP was found along the dorsal midline between the developing cerebral cortices, which were gently separated to expose the tissue and using a scalpel was cut away from developing pial membranes. After collecting the 3V ChP, a scalpel was used to perform a bilateral cut along the midline to separate the developing cortex into two hemispheres. Each hemisphere was stabilized with forceps and a third of the rostral end was cut, the developing hippocampus was rolled out using the flat surface of a scalpel and the attached LV ChP was gently separated from the hippocampus/fornix using forceps.

### FACS purification of healthy embryonic cells

Lateral, third and four ventricles of three litters of embryonic E16.5 dpc embryos were rapidly dissected and pooled together in dissecting medium (HBSS+06% Glucose+1xPen/Strep+Filtered medium 0.22 filter) (Corning, Cat:21-023-CV) + glucose (Sigma, Cat:7021-100G). Dissected ChPs were spun down at 300xG for 3 mins in a centrifuge. Whole ChP was dissociated enzymatically by preparing fresh solution of collagenase II (Gibco, 17101-015) supplemented with 3 mM calcium (Sigma, C3881) for 15 minutes at room temperature. Next, CP was tapped and flicked 30 times and incubated at 37°C incubator 3 consecutive times. The enzymatic solution with digested ChP tissue was then diluted using 1xHBSS, followed by pelleting down cells at 300xG for 3 mins. TrypLE (Life Technologies, TrypLE, Catalog: 12604) was then added and samples were incubated for 5 minutes followed by trituration using a micropipette. Samples were finally washed with ChP epithelial cell (CPEC) medium (DMEM+10%FBS+1xPen/strep) and re-suspended in 1xHBSS + glucose + LIVE/DEAD staining kit (ThermoFisher, Catalog: L-3; 224). Live cells identified by green fluorescence (calcein-AM) (LifeTechnologies, Catalog L3224A) and lack of EtD-1 homodimer staining (L3224B) were selected and sorted by a MoFlo Astrios cell sorter (Beckman Coulter) into 500 µL filled into dissecting medium.

### Single cell RNA-seq

Single cells were processed through the 10X Genomics Single Cell 3’ platform using the Chromium Single Cell 3′ Library & Gel Bead Kit V1 and V2 kit (10X Genomics), as per the manufacturer’s protocol. Briefly, 7,000 cells were loaded on each channel and partitioned into Gel Beads in Emulsion in the Chromium instrument where cell lysis and barcoding occur. This was followed by amplification, fragmentation, adaptor ligation and index library PCR. Libraries were sequenced on an Illumina HiSeqX at a read length of 98 base pairs.

### Tissue fixation and processing

Dissected ChP were fixed in 4% paraformaldehyde (in 1X phosphate-buffered saline, pH 7.4). For microtome sectioning, embryonic brains were drop fixed in 4% paraformaldehyde, and incubated in the following series of solutions in 1X phosphate-buffered saline: 10% sucrose, 20% sucrose, 30% sucrose, 1:1 mixture of 30% sucrose and Optimal Cutting Temperature (OCT) compound (overnight) and finally in OCT compound alone (1 hour). Samples were then frozen in OCT compound.

### Immunohistochemical analysis

Cryosections were permeabilized with 0.1% Triton X-100 in phosphate-buffered saline, and incubated with primary antibodies overnight and then with secondary antibodies for 2 hours. Sections were counterstained with Hoechst 33342 (Thermo Fisher Scientific) and mounted on slides using Fluoromount-G (SouthernBiotech, Birmingham, AL).

For Ki67 staining, an antigen retrieval step was included: A food steamer (catalog number 5712, Oster, Boca Raton, FL) was filled with water and preheated until the chamber temperature reached 100°C; sections were then immersed in boiling citric acid buffer (10 mmol/L sodium citrate, 0.05% Tween 20, pH 6) and placed in the steamer for 20 minutes. Sections were then cooled to room temperature.

### Antibodies and Probes

We used the following antibodies: AQP1 – mouse (1:100) – Santa Cruz – sc-32737; PECAM1 – rat (1:200) – BD Pharmingen – Cat. 550274; COL1A1 – rabbit (1:250) – Abcam – ab34710; Spi1/PU.1 – rat (1:250) – Novus Bio – MAB7124-SP; KI67 – mouse (1:50) – BD Pharmingen – 550609; Ac-Tubulin – (1:250) – mouse – Sigma; Ccdc67 – rabbit (1:500) – Proteintech – 24579-1-AP; Shisa8 – rabbit (1:500) – abcam – ab188621; Rtn1 – rabbit (1:1000) – abcam – ab83049; Tubb3 – mouse (1:250) – Biolegend – 801202; 5-HT – goat (1:3000) – Sigma Aldrich – S5545; Slc40a1 – rabbit (1:250) – Alpha Diagnostic International – MTP11-A; LYVE-1 – rat (1:300) – R and D – AF2125; Spi-C –rabbit (1:35) – ThermoFisher – PA5-67537; CLDN5-488 – mouse (1:400) – Invitrogen – Cat. 331588; VWF – rabbit (1:200) – ThermoFisher – Cat. MA5-14029; ACTA2-Cy3 – mouse (1:200); MRC1/CD206 – rabbit (1:250) – Abcam – ab64693; Erg – rabbit (1:300); GFAP – mouse (1:1000)

We used the following RNAscope probes: Ins2 (mm-Ins2-C2, Ref: 310751-C2, Lot: 18067A)

### Whole explant RNAscope *in situ* hybridization and immunohistochemistry

For whole mount smFISH, lateral, third, and fourth ventricle choroid plexus explants were dissected from E16.5 embryos and fixed with 4% paraformaldehyde (PFA) for 10 minutes at room temperature in a 9-well glass plate before beginning the manufacturer’s provided protocol for RNAscope Fluorescent Multiplex (ACD). Free-floating explants were incubated with Target Retrieval Reagent (ACD) in a vegetable steamer for 12 minutes. Subsequently, explants were washed 3×3 minutes in double distilled water prior to incubation with Protease III Reagent (ACD) for 8 minutes at 40°C, followed by another 3×3 minute wash cycle in double distilled water. Target Probes (ACD) were then hybridized and amplified according to the manufacturer’s specifications. After hybridization, immunohistochemical staining was performed in a subset of explants and described above. Allen Brain Atlas^73^ and Gene-paint^36^ were used to obtain *in situ* hybridization images of transcript localization within embryonic and adult brains.

### Scanning and transmission electron microscopy

Lateral ventricle choroid plexus tissue from embryonic and adult brain was micro-dissected in HBSS (Thermo Fisher Scientific) and drop-fixed immediately in FGP fixative (5% Glutaraldehyde, 2.5% Paraformaldehyde and 0.06% picric acid in 0.2 M sodium cacodylate buffer, pH 7.4). After 2 hour fixation at room temperature, the choroid plexus tissue was washed in 0.1M cacodylate buffer and postfixed with 1% Osmiumtetroxide (OsO4)/1.5% Potassiumferrocyanide (KFeCN6) for 1 hour, washed twice in water, and once in 50mM Maleate buffer pH 5.15 (MB) and incubated in 1% uranyl acetate in MB for 1hr followed by 1 wash in MB, 2 washes in water and subsequent dehydration in grades of alcohol (10 min each; 50%, 70%, 90%, 2×10 min 100%). The samples were then put in propyleneoxide for 1 hr and infiltrated ON in a 1:1 mixture of propyleneoxide and TAAB Epon (TAAB Laboratories Equipment Ltd, https://taab.co.uk). The following day, the samples were embedded in TAAB Epon and polymerized at 60°C for 48 hrs. Ultrathin sections (about 80nm) were cut on a Reichert Ultracut-S microtome, picked up on to copper grids stained with lead citrate. Images were acquired with a JEOL 1200EX transmission electron microscope, and recorded with an AMT 2k CCD camera (Biological Electron Microscopy Facility, Harvard Medical School).

### Whole explant viral transduction

Whole ChP were isolated and floated on Polycarbonte Track-Etched membranes (Whatman, Nucleopore, 8.0 µm Pore Size, Diameter, 13 mm) membranes in DMEM and infected with AAV-Syn1-GFP (AAV1.Syn.GCaMP6s.WPRE.SV40) (Penn Vector P2824), while incubated in DMEM for 48 hours in 37°C incubators under (95%O2/5%O2). Explants were then stained with antibodies against Tubb3 (Tuj1, Biolegend, 1:250) and with corresponding secondary antibodies to visualize neurons, while native fluorescence was used to visualize Syn1 expression. Imaging was performed with a Zeiss LSM 700 confocal laser scanning microscope.

### Hematoxylin and Eosin (H&E) staining

Paraffin-embedded brains were sectioned to a thickness of 5 microns. The sections were de-paraffinized in xylene and then re-hydrated via successive incubation in 100% ethanol, 95% ethanol and water. Sections were incubated in Gill 3 hematoxylin (Sigma Aldrich, St. Louis, MO) for 2 minutes, followed by a 5-second incubation in 0.5% ammonia water to increase the contrast of the hematoxylin stain. Next, sections were rinsed in water and incubated in 1% alcoholic eosin for 3 minutes. Finally, sections were dehydrated via successive incubation in 95% ethanol, 100% ethanol and mounted using Permount (Thermo Fisher Scientific).

### Adult ChP tissue dissection

Adult brain was first separated into hindbrain and connected mid/forebrain. From the hindbrain unit, the cerebellum was lifted to expose the cisterna magna and lateral arms of each ChP were separated using a scalpel. The 3V ChP was found in the medial space between in cerebral cortices and was exposed using a scalpel to gently separate the cortices and cut contra-lateral projections from each hemisphere. Next, irrigating this space with 1xHBSS revealed the ChP, which was clearly identifiable by a blood vessel running along its rostro-caudal axis. The 3V ChP travels ventrally as it extends into the rostral region of the brain, to connect to the ventral roots of the LV ChP in each hemisphere. The 3V ChP was separated from each of these connecting structures using scalpels. Next, to collect the LV ChP, cortices were separated into the two hemispheres by a bilateral cut along the midline. A third of each hemisphere from the rostral end was cut, and the hippocampus was rolled out using the flat surface of a scalpel and the attached LV ChP was separated from the hippocampus/fornix using a scalpel.

### Single nucleus RNA-seq

Working on ice throughout, ChP tissue was transferred into the dounce homogenizer (Sigma Cat No: D8938) with 2mL of EZ Lysis Buffer (Sigma-Aldrich: NUC101-1KT). Tissue was carefully dounced while on ice 25 times with Pestle A followed by 25 times with Pestle B, then transferred to a 15mL conical tube. 2mL of EZ lysis buffer were added to the dounce homogenizer to rinse residual nuclei and transferred to 15mL tube for a final volume of 4mL. Homogenate was incubated on ice for 10 mins and then centrifuged with a swing bucket rotor at 500g for 5 mins at 4°C. Supernatant was removed and the pellet was resuspended in 100 µl of ice cold PBS + 0.04% BSA (NEB B9000S). 40µm FlowMi cell strainers were pre-wetted with ice cold 200µl of PBS and the resuspended nuclei were gently filtered through the FlowMi into 1.5mL Eppendorf tubes. Nuclei were counted using the Nexcelom Cellometer Vision and a DAPI stain. DAPI was diluted to 2.5µg/µl in PBS and 20µl of the DAPI was pipette mixed with 20µl of the filtered nuclei suspension, then 20µl of the stained nuclei were pipetted into the cellometer cell counting chamber (Nexcelom CHT4-SD100-002). Nuclei were counted using a custom assay with dilution factor set to 2. 10,000 nuclei were the input to 10X Genomics single-cell 3’ Gene Expression v2 assay. cDNA was amplified for 12 cycles and resulting WTA measured by Qubit HS DNA (Thermo Fisher Scientific: Q32851) and quality assessed by BioAnalyzer (Agilent: 5067-4626). This WTA material was diluted to <8ng/µl and processed through v2 library construction according the manufacturer’s protocol, and resulting libraries were quantified again by Qubit and BioAnalzyer. Libraries were pooled and sequenced on 2 lanes of Illumina HiSeqX by The Broad Institute’s Genomics Platform.

## STATISTICAL ANALYSIS

### Regionalized proliferation in the LV ChP

To quantify the proliferating front of the LV ChP, tissue was stained with Ki67 and Hoechst. Immuno-stained explants were imaged using LSM 710 laser scanning confocal microscope (Zeiss) with tiling function to collect images of whole LV ChP. The tissue was divided into three equal regions along the dorso-ventral axis of each LV ChP, into brain-proximal, medial and distal regions (excluding the distalmost ventricular of the tissue). Hoechst staining was used to identify cellular nuclei and proliferating cells were identified by overlapped Ki67 staining. Percentage of proliferating cells were counted in each region. A total of four independent explants were quantified. Multiple comparisons ANOVA was employed using statistical software package Graphpad PRISM. Data are represented as means ±SEM. *P* < 0.05 was considered significant.

### Pre-processing of droplet-based scRNA-seq

De-multiplexing, alignment to the mm10 transcriptome and unique molecular identifier (UMI)-collapsing were performed using the Cellranger toolkit from 10X Genomics (version 1.1.0 for the first experiment, which was V1 chemistry, and version 2.0. for experiments two and three, which were V2 chemistry). For each cell, we quantified the number of genes for which at least one read was mapped, and then excluded all cells with fewer than 500 detected genes. Since the total number of UMI and genes detected varies across cell types (Fig. S1A), we further excluded epithelial cells with fewer than 10,000 total UMI and mesenchymal cells with fewer than 6,000 total UMIs. Genes that were detected in less than 5 cells were excluded. Expression values *E_i_*_,*j*_ for gene *i* in cell *j* were calculated by dividing UMI counts for gene *i* by the sum of the UMI counts in cell *j*, to normalize for differences in coverage, and then multiplying by 10,000 to create TPM-like values (TP10K), and finally computing log_2_(TP10K + 1). Batch correction was performed for each cell type separately using ComBat as implemented in the R package *sva*^74^, using the default parametric adjustment mode. The output was a corrected expression matrix, which was used as an input to further analysis.

### Pre-processing of droplet-based snRNA-seq data

De-multiplexing, alignment to the mm10 transcriptome and unique molecular identifier (UMI)-collapsing were performed using the Cellranger toolkit (version 2.1.1, chemistry V2) provided by 10X Genomics. For each cell, we quantified the number of genes for which at least one read was mapped, and then excluded all cells with fewer than 400 detected genes. Genes that were detected in less than 10 cells were excluded. Expression values *E_i_*_,*j*_ for gene *i* in cell *j* were calculated by dividing UMI counts for gene *i* by the sum of the UMI counts in cell *j*, to normalize for differences in coverage, and then multiplying by 10,000 to create TPM-like values (TP10K), and finally computing log_2_(TP10K + 1). For ease of data handling, and since ∼80% of cells were epithelial cells, epithelial cells from the second batch of the adult data were randomly down-sampled to 6,000 cells. To address batch effects, we used canonical correlation analysis (CCA) from the *Seurat* package^75^ on the union of the top 2,000 variable genes of batch 1 and batch 2 of all nuclei data, including embryo and adult. When aligning the CCA subspaces, we used 20 dimensions. Dimensionality reduction to 2D for visualization was performed using UMAP^76^ on the first 11 CCA dimensions. Clustering of cells was performed using the FindClusters() function from the *Seurat* package in R on the first 20 CCA dimensions, with a resolution of 0.8.

### Identifying variable genes

Selection of variable genes was performed by fitting a logistic regression to the cellular detection fraction (often referred to as α), using the total number of UMIs per cell as a predictor as in^77^. Outliers from this curve are genes that are expressed in a lower fraction of cells than would be expected given the total number of UMIs mapping to that gene, that is, likely cell-type or state-specific genes. We used a threshold of deviance between <−0.15 and <-0.3. In order to further correct for batch effect, if there were enough cells from all three batches, variable genes were calculated within each batch, and only the intersection of variable genes in each batch were taken for downstream analysis. This was the case when analyzing all cells (Fig. 1), and epithelial and mesenchymal cells. Batch was ignored for computing variable genes for immune, endothelial and neuro-/glia-like cells. We restricted the expression matrix to this subset of variable genes and values were centered and scaled and capped at a z-score of 10.

### Dimensionality reduction using PCA and visualization using t-SNE or UMAP

We restricted the expression matrix to the subsets of variable genes and high-quality cells noted above, and then centered and scaled values before inputting them into principal component analysis (PCA), implemented using ‘RunPCA’ in Seurat which runs the irlba function. The cell embeddings were either the singular vectors themselves or the singular vectors multiplied with the singular values depending on the cells. After PCA, significant principal components were identified using the elbow-method when looking at the distribution of singular values. Scores from only those significant principal components were used as the input to further analysis. For visualization purposes, the dimensionality of the datasets was further reduced to 2D embeddings using the RunTSNE() function of the *Seurat* package in R on the significant PCs.

### Clustering and removing doublets

To cluster single cells by their expression, we used an unsupervised clustering approach, based on the Infomap graph-clustering algorithm^78^. In brief, we constructed a *k*-nearest-neighbor graph on the data using, for each pair of cells, the Euclidean distance between the scores of significant principal components to identify *k* nearest neighbors. The parameter *k* was chosen to be consistent with the size of the dataset. Specifically, *k* was set to 40 for neuronal/glia-like cells and to 20 for immune cells, for which subclusters of macrophages were *post-hoc* merged together based on expression of canonical markers. The nearest-neighbor graph was computed using the function *nng* from the R package *cccd*. The *k*-nearest-neighbor graph was then used as the input to Infomap^78^, implemented using the *infomap.community* function from the *igraph* R package. For major cell types (Fig. 1C), clusters were *post-hoc* merged to six major cell populations using canonical markers for all cell types detected. Doublets would form their own clusters and were clearly identifiable by dual expression of e.g. epithelial genes and endothelial genes and were removed from further analysis and any visualizations in the paper.

### Scoring cells using signature gene sets

In order to score cells using a gene sets, such as cell cycle (Fig. 2A) or arterial / venous gene expression (Fig. S5A), we averaged over genes in the gene set after centering and scaling gene expression across cells. For cycling cells, cells with a z-score above 2 were classified as cycling cells.

### Topic modeling

LDA was computed on epithelial, mesenchymal and endothelial cells separately in the embryo, and on epithelial cells in adult. Specifically, we used the FitGoM() function from the *CountClust* R package (Dey et al., 2017) to fit LDA topic models to the UMI counts (Bielecki et al., 2018). To improve topic signals, genes expressed in more than 98% of cells or less than 2% of cells were removed from the count matrix prior to fitting the topic models (analogous to the removal of highly abundant words or extremely rare words in document analysis), except for endothelial cells. The number of topics to fit (K) and the tolerance value are required to run FitGoM() function. Thus, for each cell type and developmental status, we fit a range of K and tolerance values and picked values that gave us robust topics and where we mostly saturated the number of informative topics found. For the embryo data, this was achieved with a tolerance of 0.1, and a K of 16 for endothelial cells, K of 26 for epithelial and mesenchymal cells, and K of 14 and tolerance of 0.01 for adult epithelial nuclei. The top genes to highlight for each topic were selected using the ExtractTopFeatures() function.

### Defining cell-type or cluster signatures

Differential expression between cell types in scRNA-Seq of Embryo (Fig. 1) or snRNA-Seq of adult (Fig. S8), and between clusters of immune cells and neuro-/glia-like cells was performed using MAST^79^, which fits a hurdle model to the expression of each gene, consisting of logistic regression for the zero component (*i.e.* whether the gene is expressed) and linear regression for the continuous component (*i.e.* the expression level). The regression model includes terms to capture the effects of the cell subset or cluster, while controlling for cell complexity (*i.e.* the total number of unique molecular identifiers (nUMI)). Specifically, we used the regression formula, ***Yi ∼ X + N***, where ***Yi*** is the standardized log_2_(TP10K+1) expression vector for gene *i* across all cells, ***X*** is a factor variable reflecting cell subset or cluster membership, ***N*** is the scaled nUMI in each cell. In all cases, the discrete and continuous coefficients of the model were retrieved and p-values were calculated using the likelihood ratio test in MAST. Q-values were estimated using the Benjamini-Hochberg correction. In order to identify cell-type specific or cluster specific markers, a FDR cutoff for the hurdle model was chosen using the elbow-method and a small mastfc cutoff to exclude genes with very small effect sizes. If a gene passed the FDR and the mastfc cutoff in only one cluster / cell type, it was considered to be specific.

### Diffusion map

For macrophages, cells of replicate 3 were excluded for this analysis because they had very high expression of mitochondrial genes. Diffusion components were calculated on a gene expression matrix limited to variable genes within the macrophages (not correcting for batch). Diffusion components were calculated using the DiffusionMap function from the *destiny* package in R (Angerer et al., 2016) with a k of 20 and a local sigma.

For diffusion analysis of epithelial and neuronal cells, we selected cells from only 3V ChP, where most neuronal cells were sampled from, only from replicate 3, because of strong batch effects between replicates. Furthermore, the epithelial cell cluster scoring highly for IEG genes (Fig. S2D) and Oligodendrocyte precursor cells were also excluded from this analysis. Of the remaining cells, variable genes were computed as described above. Diffusion components were calculated as described for macrophages with a k of 30 and a local sigma.

### Cell-cell interactions

First, a gene set table was created where cell type markers from Fig. 1C were considered to be general markers, and then subtype specific gene sets were added. For neuronal /glia-like cells and immune cells, cluster specific genes were added. For the endothelial, epithelial and mesenchymal cells, the top 50 scoring feature genes from the topics used in this paper were added. Gene names were then converted to their human homologs. We used the data in https://baderlab.org/CellCellInteractions/ as a source of cognate ligand-receptor interactions. First, we identified ligand-receptor pairs between any two gene sets. Then, between two given gene sets, we normalized the weights per ligand and then per receptor to 1, as there is redundancy. We then added up all the weights between two gene sets. In order to assess the significance for the strength of two gene sets interacting with each other, we shuffled the gene to gene-set assignment 10^4^ times, and each time recomputed the overall interaction strength between any two gene sets. This formed the null distribution against which the empirical p-value was computed.

### Cell type classification across embryo or adult

To compare the embryo scRNA-seq dataset to the adult snRNA-seq dataset, a random forest classifier was used from the R package ‘randomForest’. First, common variable genes were computed between the two datasets by taking the intersection of the variable genes in each dataset. Then, the classifier was trained on the scaled expression matrix of the common variable genes in the adult snRNA-seq dataset. The out-of-bag error was 0.15. The classifier then ran on the scaled expression matrix of the common variable genes in the embryo scRNA-seq dataset. We then compare the predicted cell type assignment of the random forest classifier to our own cell type assignment shown in Fig. 1C.

## DATA AVALABILITY

scRNA-seq and snRNA-seq data will be deposited in GEO and the Single Cell Portal (URL). Accession numbers have been requested.

## ACKNOWLEDGEMENTS

We thank members of the Lehtinen, Regev, Andermann, Fleming, Hla, and Stevens labs, and L. Tsai for reagents and helpful discussions; M. Shannon, and H. Zucker for experimental assistance and advice; M. Ericsson and HMS Electron Microscopy Facility; the Flow Cytometry facility at the Broad Institute; and A. Hupalowska and L. Gaffney with illustrations. This work was supported by: Glenn/AFAR Postdoctoral Fellowship for Translational Research on Aging and Reagan Sloane Shanley Research Internship (N.D.); EMBO Fellowship and the Helen Hay Whitney Foundation Fellowship (N.H.); NSF Graduate Research Fellowship (F.B.S.); NIH T32 HL110852 (J.C); Boston Children’s Hospital-Broad Institute Collaboration Grant (A.R. and M.K.L.); Pediatric Hydrocephalus Foundation and NIH R01 NS088566 (M.K.L.); the New York Stem Cell Foundation (M.K.L. and F.Z.); BCH IDDRC 1U54HD090255; and the Klarman Cell Observatory. F.Z. and A.R. are Investigators of the Howard Hughes Medical Institute. F.Z. and M.K.L. are New York Stem Cell Foundation – Robertson Investigators.

## AUTHOR CONTRIBUTIONS

N.D., R.H.H., N.H., O.R., A.R., and M.K.L. designed the study; N.D., J.H. and A.J. performed tissue dissections; N.D. and J.H. developed whole tissue explant smFISH and imaging protocols; N.D., D.D., C.M., L.N. and C.R. prepared libraries and performed sequencing; R.H.H. and N.H. performed all computational analysis of scRNA-seq and snRNA-seq data in the lab of A.R., with help from S.J.R.; J.C. performed EM studies; N.D. and F.B.S. performed data analysis, N.D., R.H.H., N.H., A.R. and M.K.L. wrote the manuscript, and all co-authors edited it.

